# Elevated mitochondrial metabolism in Down syndrome iPSCs reduces commitment to neuroectoderm

**DOI:** 10.1101/2025.05.09.653211

**Authors:** Malay Chaklader, Guilherme Henrique Souza Bomfim, Neil Jeju, Yingli Duan, Brian F Niemeyer, Joaquin M Espinosa, Edwin Rosado-Olivieri, Weichun Lin, Rodrigo S. Lacruz, Beverly Rothermel

**Affiliations:** Department of Internal Medicine, Division of Cardiology, University of Texas Southwestern Medical Center, 5323 Harry Hines Blvd, Dallas, TX 75390, USA; Department of Molecular Biology, University of Texas Southwestern Medical Center, 5323 Harry Hines Blvd, Dallas, TX 75390, USA; Department of Molecular Pathobiology, New York University College of Dentistry, 345 East 24^th^ St., New York, NY, 10010, USA; Linda Crnic Institute for Down Syndrome, University of Colorado Anschutz Medical Campus, 12700 East 19^th^ Avenue, Aurora, CO 80045, USA; Department of Neuroscience, Division of Cardiology, University of Texas Southwestern Medical Center, 5323 Harry Hines Blvd, Dallas, TX 75390, USA

**Keywords:** Cell cycle, Down syndrome, Mitochondrial calcium uniporter, primary cilium, Neuroectoderm

## Abstract

A key feature of Down syndrome (DS) is reduced neurogenesis. Here, we provide evidence that increased mitochondrial metabolism in DS stem cells reduces their ability to commit to neuroectoderm (NE), one of the earliest steps in the development of the nervous system. We show that mitochondria in induced pluripotent stem cells derived from individuals with DS (3S-iPSCs) have a higher membrane potential and increased capacity for calcium uptake via the mitochondrial calcium uniporter (MCU) compared to isogenic, euploid controls. Consequently, 3S-iPSCs proliferate faster and spend less time in G_1_ of the cell cycle. This reduces the opportunity for growth of a primary cilium, an important developmental signaling hub. Inhibiting MCU or slowing proliferation of 3S-iPSCs is sufficient to increase ciliation and improve commitment to NE. In summary, we provide evidence that a mitochondria-to-cilia signaling axis important during the earliest steps of neurogenesis is dysregulated in DS, yet remains amenable to small molecule intervention.

## INTRODUCTION

Down syndrome (DS) is the most common genetic cause of intellectual disability, occurring at a frequency of around 1 out of every 700 live births (Bull, 2020). Although it has been known for more than sixty years that inheriting a full or partial extra copy of human chromosome 21 (HSA21) causes DS (Lejeune *et al*, 1959), the specific molecular mechanisms responsible for the cognitive differences have remained elusive and are likely multifactorial. A key feature of DS is an overall reduction in brain size and neuronal density, notably in regions involved in executive function and memory (Klein & Haydar, 2022; Russo *et al*, 2024), thus, it is believed that impaired or delayed neurogenesis contributes to intellectual deficits. Compared to age-matched controls, cerebellar volume is reduced in fetuses with DS by 15 weeks of gestation (Guihard-Costa *et al*, 2006), however, neither the cause of this divergence or how early it occurs are known.

Induced pluripotent stem cells (iPSCs) derived from individuals with DS have been used extensively to model diverse aspects of neuronal development in DS. Neuronal progenitor cells (NPCs) derived from DS iPSCs proliferate more slowly than those derived from normal euploid iPSCs (Murray *et al*, 2015; Tang *et al*, 2021), whereas, glial, astrocyte progenitor cells (APCs) derived from DS iPSCs proliferate more rapidly (Kawatani *et al*, 2021). These altered rates of proliferation are consistent with observations in humans with DS. Surprisingly, few studies have examined the earliest steps of lineage commitment, so it is not known at what point DS iPSCs first show evidence of reduced neurogenesis or what the specific mechanisms are that contribute to this deficit.

The primary cilium is a non-motile, antenna-like, microtubular-based structure projecting from the surface of most mammalian cells. It acts as a signaling hub that regulates diverse cellular signaling pathways (Anvarian *et al*, 2019), including many of those involved in neuronal and cortical development (Karalis *et al*, 2022). Ciliopathies are a group of disorders caused by mutations in essential cilia-related genes (Waters & Beales, 2011). Similar to DS, ciliopathies display multisystemic pathologies, including brain malformations and cognitive deficits. This overlap in clinical features has been noted by several investigators who have proposed DS as a type of ciliopathy (Jewett *et al*, 2023; Raveau *et al*, 2017). The primary cilium is formed during the G_1_ phase of the cell cycle and thus is present both on cells in G_1_ and on those in G_0_ that have withdrawn from the cell cycle. The cilium is anchored at its base by a centromere and must be disassembled upon entry into S phase, to allow the centromere to replicate and carry out its mitotic functions. When human embryonic stem cells (hESCs) are allowed to differentiate spontaneously, the cells within the population that commit to neuroectoderm (NE), one of the earliest steps in the development of the nervous system, are those that proliferate more slowly and have a primary cilium (Jang *et al*, 2016).

Mitochondria are not static but undergo a constant process of fission and fusion that in part helps to regulate their metabolic activity. Specific changes in mitochondrial dynamics are associated with progression through the cell cycle (Colpman *et al*, 2023; Spurlock *et al*, 2020). In proliferating cells mitochondrial fusion occurs at the G_1_ to S transition (Mitra *et al*, 2009; Schieke *et al*, 2008). This increases mitochondrial membrane potential (*ΔΨmt*) and the capacity for mitochondria to take up calcium (Ca^2+^) through the mitochondrial Ca^2+^ uniporter (MCU), which is *ΔΨmt-*dependent (Garcia *et al*, 2023; Koval *et al*, 2019). Uptake of Ca^2+^ promotes progression through the cell cycle by activating enzymes in the TCA cycle to increase ATP production (Zhao & Pan, 2021). Consequently, inhibiting MCU can reduce the rate of cell proliferation (Garcia *et al*., 2023; Koval *et al*., 2019).

We previously showed that mitochondria are more metabolically active in iPSCs derived from individuals with DS (3S-iPSCs) compared to mitochondria in isogenic, euploid control iPSCs (2S-iPSCs) (Parra *et al*, 2018). Here, we provide evidence of a causative link between this abnormally high mitochondrial activity and a reduced ability to commit to a neurogenic lineage. Mechanistically, we show that 3S-iPSCs proliferate more rapidly, thereby reducing the opportunity for growth of the primary cilium that is required for commitment to NE. Furthermore, inhibiting either cell cycle progression or MCU activity in 3S-iPSCs is sufficient to increase their neurogenic potential.

## RESULTS

### Commitment to NE is reduced in DS iPSCs compared to isogenic, euploid controls

Reduced neurogenesis is a prominent feature of DS, both during development and across the lifespan (Klein & Haydar, 2022), however, how early this deficiency first manifests is unknown. To determine whether the capacity for neurogenesis is impacted at the earliest stages of lineage commitment, we assessed the ability of 3S-iPSCs, trisomic for human chromosome 21 (Hsa21), to commit to NE under either directed or spontaneous differentiation protocols, comparing their responses to those of isogenic, 2S-iPSC controls. Prior to differentiation, both genotypes showed uniform expression of pluripotency markers as we have previously reported **(Fig. EV1A-EV1C)** (Parra *et al*., 2018). To test directed differentiation, iPSCs were shifted from mTESR-Plus^®^ maintenance media to STEMdiff™ Trilineage Ectoderm Differentiation media **(Fig. 1A)**. Five days later, transcript and protein levels of the NE lineage marker, *Paired Box 6* (*PAX6*), were assessed (Kuang *et al*, 2019). Immunohistochemistry using an antibody against PAX6 showed an abundance of PAX6-positive (PAX6+) cells in the cultures differentiated from 2S-iPSCs, whereas the frequency of PAX6+ cells in the cultures derived from 3S-iPSCs was significantly less **(Fig. 1B)**. Consistent with this, *PAX6* transcript levels, normalized to *HPRT* transcript levels, increased over a 100-fold in the 2S-derived cultures, whereas the increase in *PAX6* transcript levels in the 3S-derived cultures was significantly less **(Fig. 1C)**. Flow cytometry was used to compare the intensity and distribution of PAX6+ cells on a population level **(Fig. EV2A-EV2C)**. The median fluorescence intensity of PAX6+ signal was significantly lower in the 3S-derived cultures when compared to the PAX6+ signal in the 2S controls (**Fig. 1D and 1E**). Taken together these findings suggest a reduction, or delay, in the commitment of 3S-iPSCs to NE under directed conditions.

**Figure 1:**
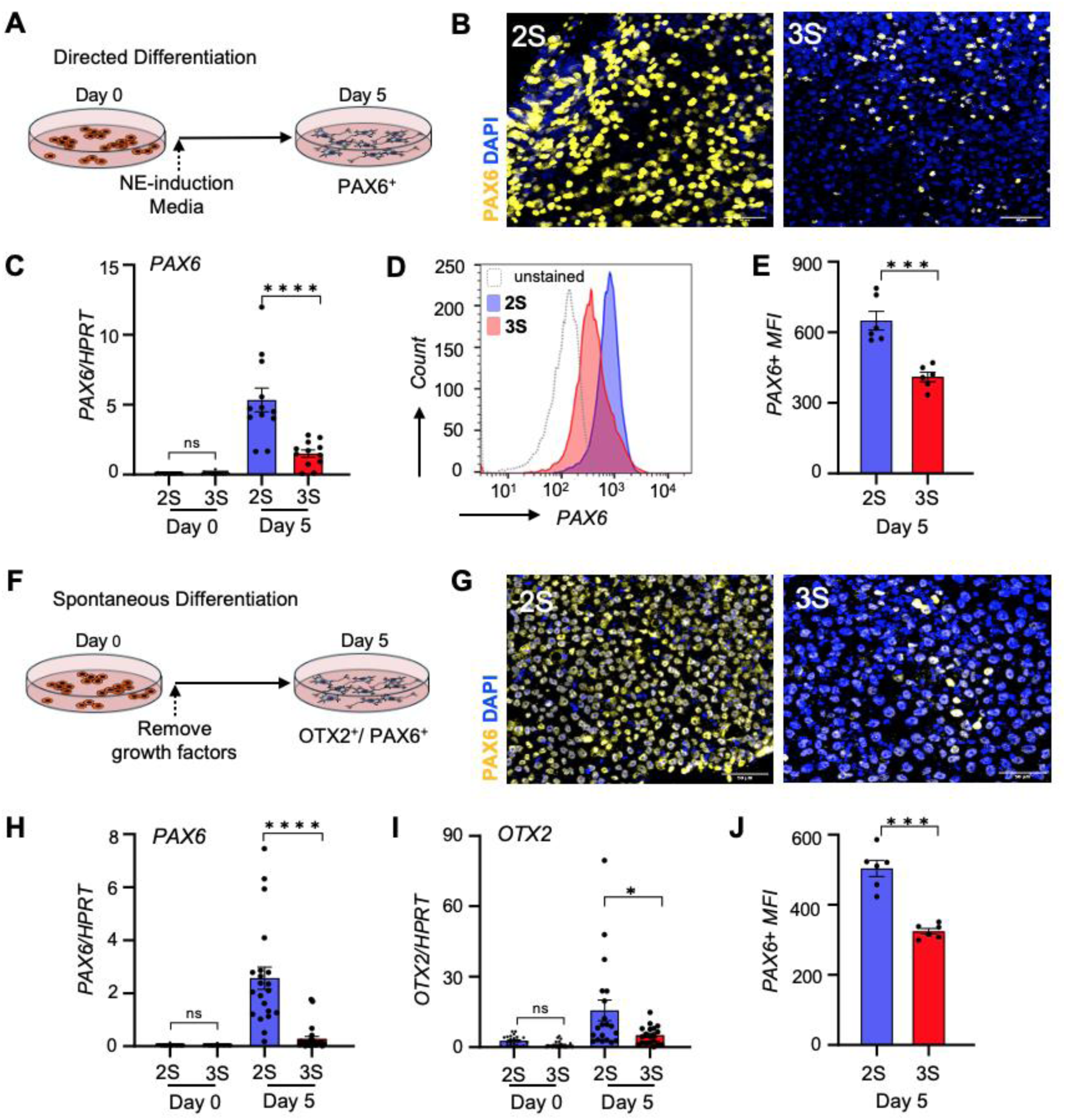
Commitment of 3S-iPSCs to Neuroectoderm is reduced compared to isogenic, 2S-iPSCs. **(A)** Schematic for directed differentiation of iPSCs to neuroectoderm (NE) (used in Fig 1B-1E). **(B)** Fewer cells are PAX6-positive (PAX6+) (yellow) in 3S-derived cultures than in 2S-derived cultures after 5 days of ectoderm-directed differentiation. DAPI-stained nuclei are in blue (Scale bar = 50μm). **(C)** *PAX6* transcript levels are lower (n=12) and **(D and E)** PAX6 medial fluorescence intensity (*PAX6+-MFI*) assessed by flow cytometry is lower (n=6) in 3S-derived cultures than in 2S-derived cultures on day 5 of directed differentiation. **(F)** Schematic for spontaneous differentiation of iPSCs (used in Fig. 1G-1J). **(G)** Fewer cells are PAX6+ by immunohistochemistry (yellow) in 3S-derived cultures at day 5 of spontaneous differentiation (Scale bar = 50μm). *PAX6* **(H)** and *OTX2* **(I)** transcript levels are lower in the 3S-derived cultures on day 5 (n=18). **(J)** PAX6+ MFI is lower in 3S-derived cultures on day 5 (n=6). *Transcript levels are normalized to HPRT. Comparisons by ANOVA or Two-tailed T-test. Error bars represent SEM*. *\* P≤0.05, ** P ≤0.01, *** P ≤0.001, **** P ≤0.0001*

To compare lineage commitment during spontaneous differentiation, growth factors were removed from the media, releasing the cells to differentiate spontaneously (**Fig. 1F**). The frequency of cells staining positive for PAX6+ after 5 days of differentiation was significantly less in the trisomic 3S-derived cultures than in the isogenic 2S-derived cultures (**Fig. 1G).** The frequency of cells staining positive for Orthodenticle Homeobox 2 (OTX2+), another NE marker, was also reduced in the 3S-derived cultures **(Fig. EV2D**). Transcript levels for both *PAX6* and *OTX2* were significantly lower in the 3S-derived cultures (**Fig. 1H and 1I)** as was PAX6+ median fluorescence intensity (**Figs. 1J and EV2E)**. This suggests that a cell-autonomous factor contributes to reduced neurogenesis in DS and impacts the earliest stages of lineage commitment.

### DS 3S-iPSCs proliferate more rapidly than 2S-iPSCs. and treatment with a cell cycle inhibitor increases their commitment to NE

We noticed that our trisomic lines needed to be passaged more frequently than their isogenic, euploid controls suggesting a shorter cell cycle. Tracking changes in cell density over time verified that the rate of growth was faster in 3S-iPSC cultures than in 2S-iPSC cultures (**Fig. 2A**). Flow cytometric analysis of propidium iodide-stained, log phase growth cultures revealed that a greater percentage of 3S-iPSCs were in S phase (p=0.0002) and a lower percentage were in G_1_/G_0_ (p=0.003) compared to 2S-iPSC controls (**Fig. 2B and 2C**), consistent with the 3S-iPSCs having a shorter cell cycle. A short G_1_ phase is a characteristic feature of embryonic stem cells (ESCs) (Becker *et al*, 2006), which then must slow their rate of proliferation to initiate differentiation (White & Dalton, 2005). Lengthening of G_1_ in neural progenitors has been found to be both necessary and sufficient to trigger commitment to neurogenesis (Calegari & Huttner, 2003; Lange & Calegari, 2010; Lange *et al*, 2009). We tested whether pretreating log-phase growth 3S-iPSCs with a cell cycle inhibitor to increase the fraction of cells in G_1_ would improve the ability of 3S-iPSCs to commit to NE. Cultures were pretreated for 16 hours with either the cyclin-dependent kinase inhibitor PD-0332991 (PD) or vehicle control **(Fig. 2D and 2E)**. PD pretreatment decreased the percentage of cells in S phase and increased the percentage in G_1_/G_0_ in both the 2S and 3S iPSCs (**Fig. 2F**). PD was then removed and the cultures shifted to media that allowed spontaneous differentiation. Following 5 days of differentiation both *PAX6* transcript levels (**Fig. 2G)** and PAX6+ median fluorescence intensity (**Fig. 2H)** were significantly higher in the PD-treated cultures of both backgrounds compared to vehicle-treated controls (**Fig. 2G and 2H**). Of note, following PD treatment, there was no longer a significant difference in the level of *PAX6* expression between the two genotypes, suggesting that the trisomic and euploid iPSCs may have a similar maximal potential for commitmenting to NE, but that the more rapid rate of 3S-iPSC proliferation is a contributing factor in their reduced or delayed commitment to NE.

**Figure 2:**
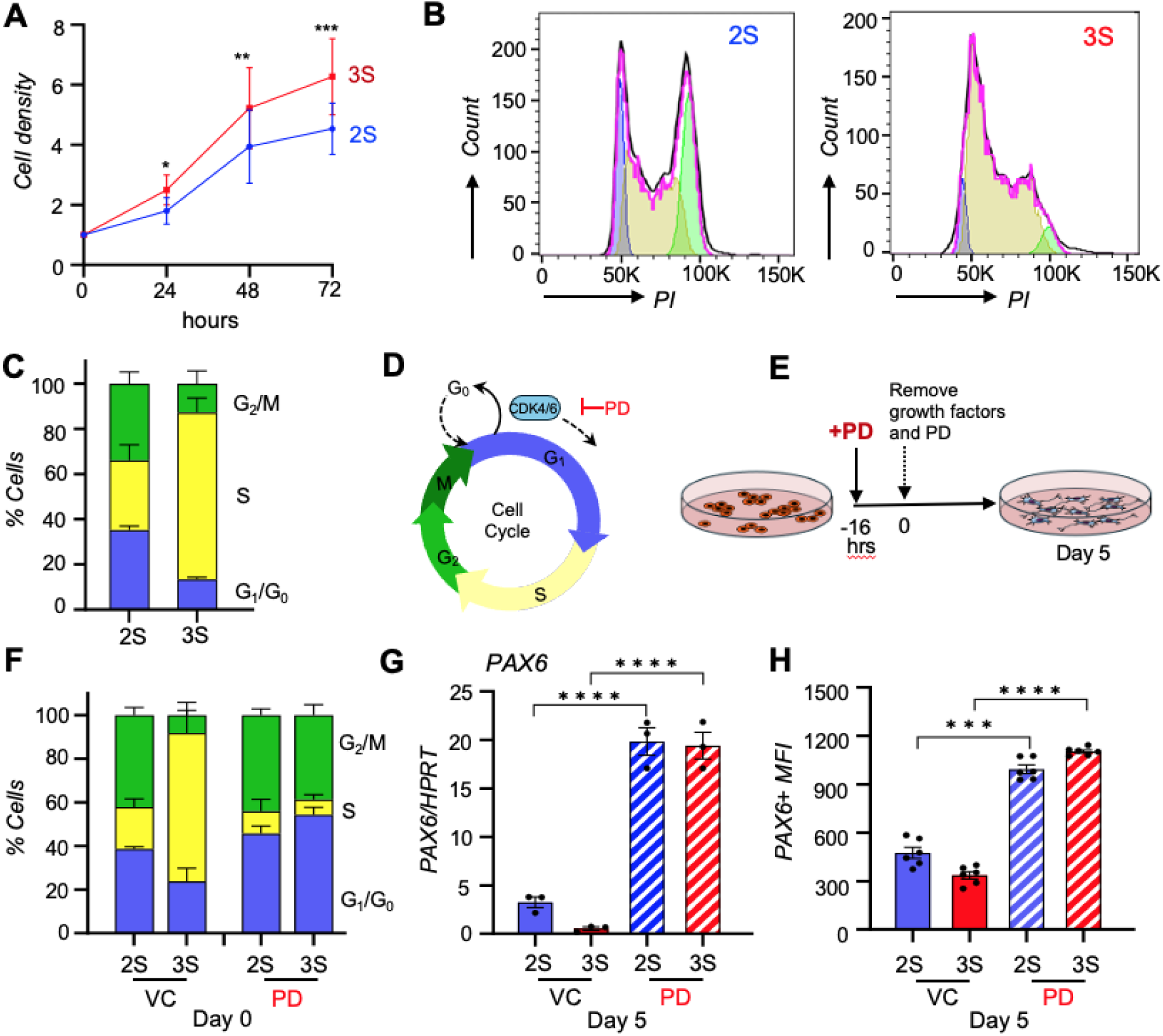
3S-iPSCs have a shorter cell cycle than 2S-iPSCs and treatment with a cell cycle inhibitor increases their commitment to NE. **(A**) Tracking changes in cell density over time shows that the 3S-iPSC cultures proliferate more quickly than 2S-iPSC cultures. **(B)** Representative flow analysis of propidium Iodide (PI) stained cultures in log-phase growth. Estimate of G_1_/G_0_ population is depicted in blue, G_2_ is in green, and S phase in yellow. **(C)** Quantification of cell cycle analysis shows that 3S-iPSC cultures contain significantly fewer cells in G_1_/G_0_ of the cell cycle and more in S phase when compared to 2S-iPSC cultures (n=3 experiments, in duplicate). **(D)** The CDK4/CDK6 inhibitor PD0332991 (PD) arrests cell cycle in G_1_. **(E)** Schematic for pretreatment with PD (5µM) or vehicle 16 hours prior to removal of growth factors to initiate spontaneous differentiation (used in Fig 2F-2H). **(F)** Pretreatment with PD increases the fraction of 3S-iPSCs in G_1_/G_0_ and decreases the fraction in S phase, consistent with a lengthening of the cell cycle. **(G)** Pretreatment with PD increases *PAX6* transcript levels and **(H)** and PAX6+ median fluorescence intensity at 5 days of differentiation (n=6). *Transcript levels are normalized to HPRT. Comparisons by ANOVA or Two-tailed T-test. Error bars represent SEM*. *\* P≤0.05, ** P ≤0.01, *** P ≤0.001, **** P ≤0.0001*.

### Primary ciliation is reduced in Down syndrome 3S-iPSCs, and depleting primary cilia in 2S-iPSCs reduces their ability to commit to NE

The primary cilium serves as a hub for a diversity of signaling pathways (Gopalakrishnan *et al*, 2023). It is present during the G_1_/G_0_ phases of the cell cycle but must be disassembled to re-enter the cell cycle and allow replication of centrosomes (**Fig. 3A**). It’s been observed that when human ESCs (hESCs) are released to differentiate spontaneously, the cells within the population that become NE are those with longer cell cycles that develop cilia (Jang *et al*., 2016). Immunofluorescence microscopy was used to compare ciliation in the 2S and 3S cultures two days after release to spontaneous differentiation. An antibody against ADP-ribosylation factor-like protein 13B (ARL13B) was used to detect cilia, whereas pericentrin (PCNT) was used to detect centrosomes, which are present in all cells but also mark the base of the primary cilium when one is present. Thus, convergent staining for both ARL13B and PCNT was used to identify ciliated cells (**Fig. 3B**). After 48 hours of spontaneous differentiation, approximately 30% of the cells in the 2S cultures were positive for both PCNT and ARL13B, indicating the presence of a primary cilium, whereas cilia-positive cells were rare in the 3S-derived, trisomic cultures (**Fig. 3C and 3D**). Depleting primary cilia has been shown to reduce the ability of hESCs to commit to NE (Jang *et al*., 2016). To verify that the primary cilium is also important for commitment of iPSCs to NE, we used siRNA knockdown of either *Kinesin Family Member 3A* (*KIF3A*) or *Intraflagellar Transport 20 (IFT20*) to reduce cilia formation in normal, euploid 2S-iPSCs. Two days after transfection with siRNA cultures were shifted to spontaneous differentiation media (**Fig. 3E**). Transcript knockdown was confirmed by qRT-PCR at day 0 (**Fig. EV3A and EV3B)** and was maintained after five days of differentiation (**Figure 3F**). Knockdown of either *KIF3A* or *IFT20* significantly reduced the percentage of 2S cells with primary cilia on day 2 of differentiation (**Fig. 3G and 3H**). It also significantly reduced *PAX6* transcript levels on day 5 of differentiation (**Figure 3I)** as well as PAX6+ median fluorescence intensity (**Fig. 3J)**, confirming the importance of the primary cilium in the commitment of iPSCs to NE. We then asked whether pretreatment of 3S-iPSCs with PD, as in Figure 2, was sufficient to increase the frequency of ciliated cells, and indeed this was the case (**Fig. EV3C and EV3D)** placing cell cycle length upstream of primary cilia formation in the progression to NE lineage commitment.

**Figure 3:**
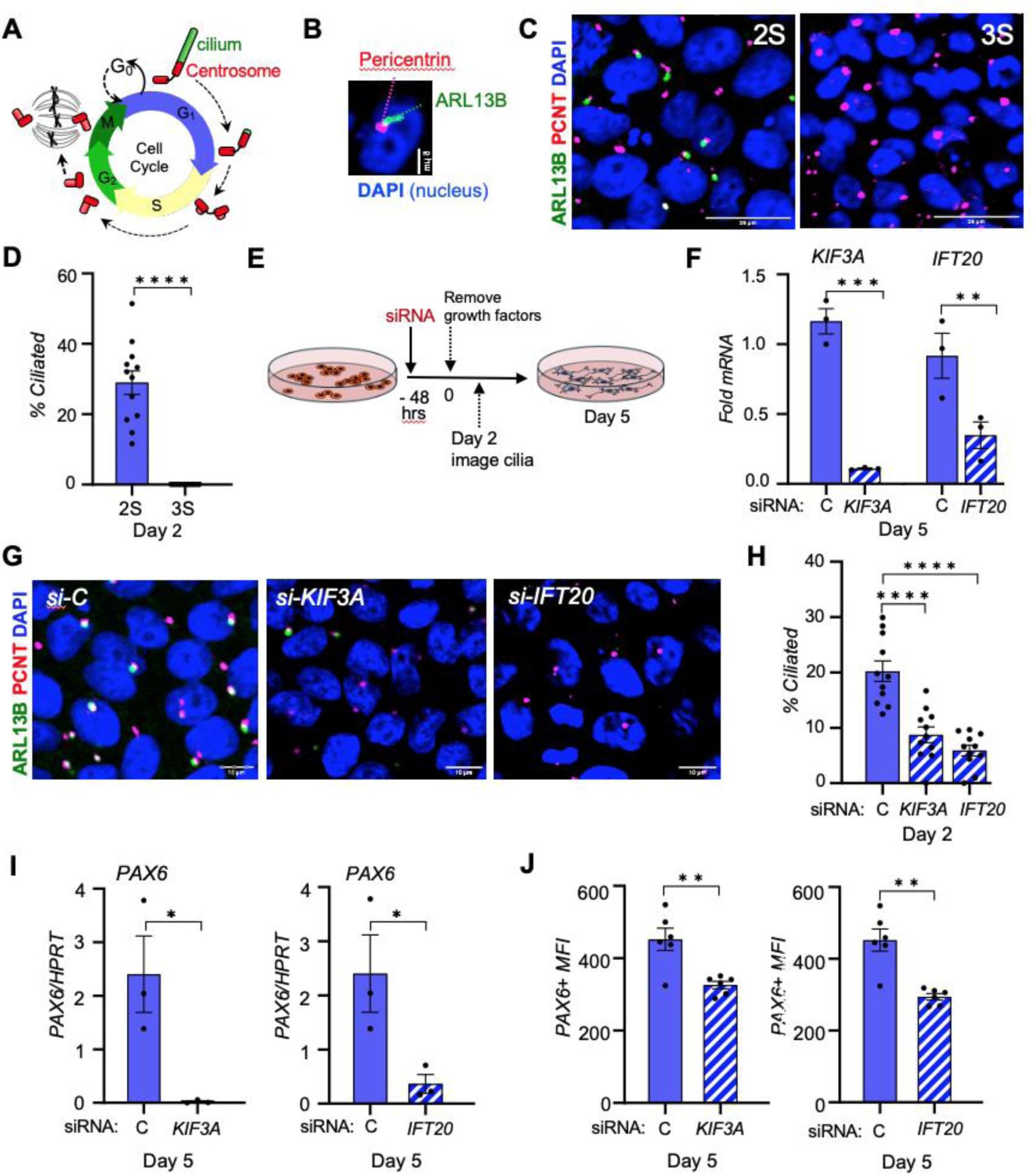
Primary ciliation of 3S-iPSCs is reduced. **(A)** Schematic depicts changes in the primary cilium and centrosomes across the phases of the cell cycle. **(B)** Co-staining with antibodies against ARL13B (green) and PCTN (red) was used to identify cells with primary cilia. DAPI-stained nuclei are in blue **(C)** Representative images show immunohistochemistry for ARL13B and PCTN in 2S and 3S-derived cultures on day 2 of spontaneous differentiation (Scale bar = 25μm). **(D)** The fraction of cells with primary cilia is significantly lower in the 3S-derived cultures (n=3, multiple fields each). **(E)** Schematic depicts experimental timeline for siRNA knockdown and differentiation. **(F)** SiRNA depletion of *KIF3A* and *IFT20* transcripts in the 2S-derived cultures is sustained over 5 days of spontaneous differentiation compared to a non-specific control siRNA (n=3). **(G)** Representative images show ARL13B and PCTN co-staining on day 2 of spontaneous differentiation of 2S-iPSCs transfected with the indicated siRNAs (Scale bar = 10μm). **(H)** The fraction of cells with primary cilia is reduced in the 2S cultures depleted for either *KIF3A* or *IFT20* (n= 3, multiple fields). **(I)** *PAX6* transcript levels are reduced in the 2S cultures depleted of either *KIF3A* or *IFT20* on day 5 of differentiation (n=3). **(J)** Knockdown in 2S-iPSCs reduces PAX6+ median fluorescence intensity at 5 days of spontaneous differentiation (n=6). *Transcript levels are normalized to HPRT. Comparisons by ANOVA or Two-tailed T-test. Error bars represent SEM*. *\* P≤0.05, ** P ≤0.01, *** P ≤0.001, **** P ≤0.0001*.

### Inhibiting MCU activity increases G_1_/G_0_, ciliation, and commitment to NE of 3S-iPSCs

Specific changes in mitochondrial dynamics are associated with progression through the cell cycle (Colpman *et al*., 2023; Spurlock *et al*., 2020). In proliferating cells, mitochondria fusion occurs at the G_1_/S transition, thereby increasing mitochondrial membrane potential (*ΔΨm*) and the production of ATP to support loading of replication factors on to DNA and subsequent DNA synthesis (Mitra *et al*., 2009; Schieke *et al*., 2008) (**Fig. E4A)**. We previously reported that the mitochondrial network is hyperfused and more metabolically active in 3S-iPSCs than in isogenic, 2S-iPSCs (Parra *et al*., 2018). Consistent with this, *ΔΨm* and mitochondrial superoxide production were elevated in 3S-iPSCs compared to in 2S-iPSCs (**Fig. EV4A and EV4B)**. The capacity for mitochondrial uptake of Ca^2+^ via MCU, which is *ΔΨm*-dependent, was also higher in permeabilized 3S-iPSCs compared to controls **(Fig. 4A-4C and Fig. EV4D-E**). The MCU inhibitor RuRED was used to verify that the increase in [Ca^2+^]_mt_ was MCU-dependent. Increased Ca^2+^ in the mitochondrial matrix increases ATP production by activate TCA cycle enzymes (De Mario *et al*, 2023). Consequently, inhibiting MCU will slow cycle progression at the G_1_/S transition, increasing the population of cells in G_1_ (Koval *et al*., 2019). To test whether inhibiting MCU activity could increase commitment of 3S-iPSCs to NE, cultures were pretreated for 16 hours with either vehicle, or the cell-permeable MCU inhibitor, Ru265, prior to releasing the cells for spontaneous differentiation. Pretreatment of 3S-iPSCs with 10 nM Ru265 for 16 hours significantly increased the fraction of cells in G_1_/G_0_ *(p<0.0001)* (**Fig. 4D**) as well as the percent of 3S-derived cells with primary cilia on day 2 of spontaneous differentiation (**Fig. 4E and 4F**). On day 5 of spontaneous differentiation, both *PAX6* transcript levels (**Fig. EV4F)** and PAX6+ median fluorescence intensity (**Fig. 4G**) were also higher in the 3S-derived cultures pretreated with Ru265 compared to vehicle treated. Similarly, siRNA-mediated knockdown of MCU transcripts 48 hours prior to differentiation (**Fig. EV4G)** increased the percent of 3S-iPSCs in G_1_/G_0_ (**Fig. 4H**) (*p<0.0001)*, the percent of cells with primary cilia on day 2 of differentiation (**Fig. 4I and 4J**), and PAX6+ median fluorescence intensity in both 2S and 3S-derived cultures on day 5 of differentiation (**Fig. 4K**). Taken together, these data suggest that increased MCU capacity in 3S-iPSCs supports an increase in the rate of cell proliferation thereby reducing the opportunity for developing a primary cilium that is required to coordinate the signals that mediate commitment to NE.

**Figure 4:**
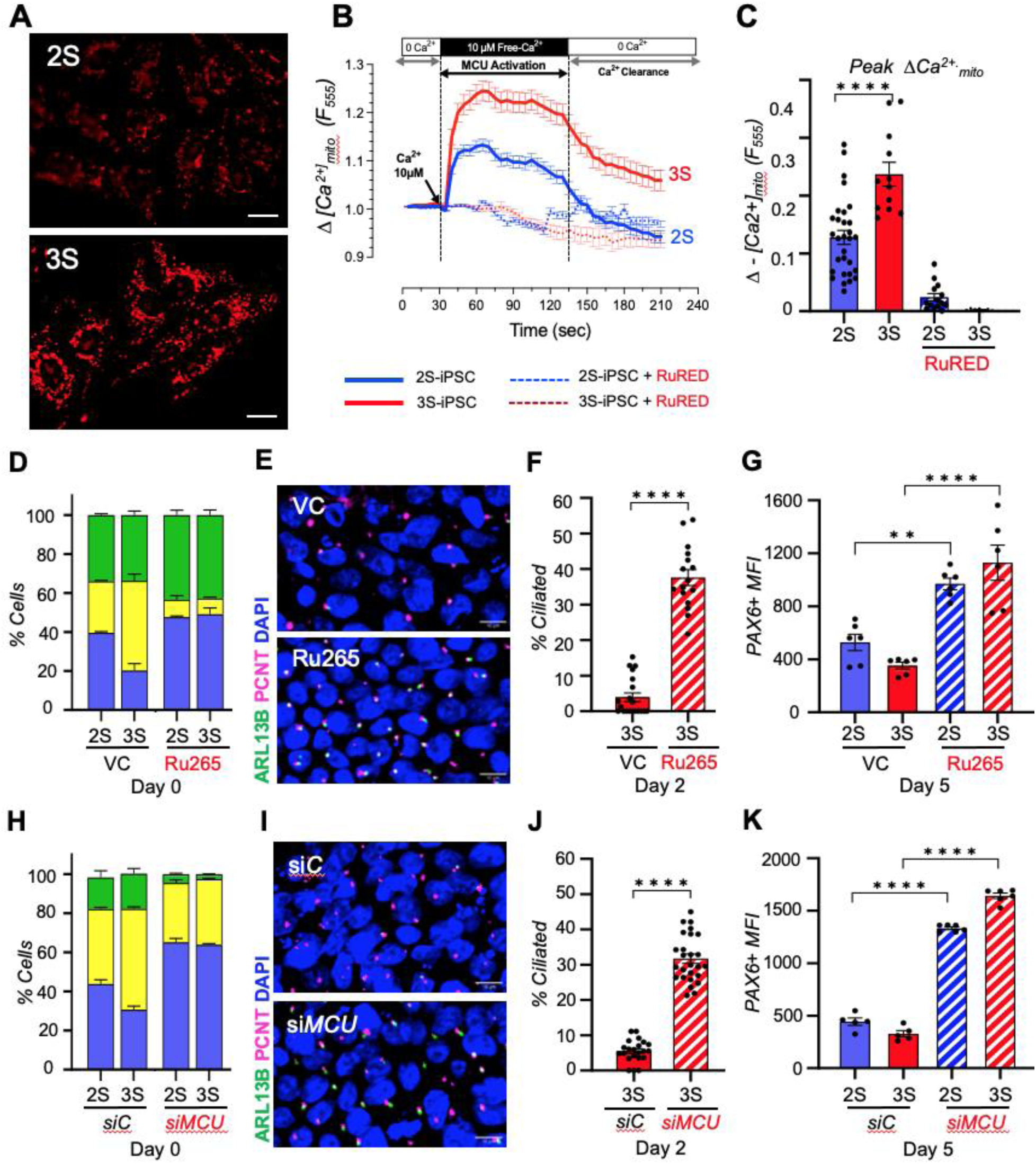
Inhibiting MCU increases commitment of 3S-iPSCs to NE. **(A)** Representative images of Rhod-2-AM signal in permeabilized iPSCs 50 seconds after the addition of 10μM free Ca^2+^ indicates enhanced mitochondrial Ca^2+^ uptake in 3S-iPSCs compared to 2S-iPSCs. **(B)** provides a temporal schematic of the protocol to assess mitochondrial Ca^2+^ uptake via MCU. The MCU inhibitor, RuRED (10μM), was used to validate MCU as the mechanism of Ca^2+^ uptake. **(C)** Peak mitochondrial uptake of Ca^2+^ is greater in 3S-iPSCs than in 2S-iPSCs. The addition of the MCU inhibitor, RuRED, was used to validate dependence on MCU. (n=3 experiments, multiple cells). **(D)** 16 hours pretreatment with 10nM Ru265 to inhibit MCU increases the fraction of 3S-iPSCs in G_1_/G_0_ (n=3, in duplicate) compared to vehicle control (VC). **(E)** Representative images show ARL13B and PCNT immunohistochemistry on day 2 of spontaneous differentiation of 3S-iPSCs pretreated with Ru265 or vehicle control (VC) (Scale bar = 10μm). **(F)** Pretreatment of 3S-iPSCs with Ru265 increases the % of cells with primary cilia on day 2 of spontaneous differentiation (n=3, 10-16 fields). **(G)** Ru265 pretreatment increases PAX6+ medial fluorescence intensity on day 5 of differentiation (n=6). **(H)** SiRNA-depletion of MCU in 3S-iPSCs for 48 hours increases the fraction of cells in G_1_/G_0_ compared to transfection with a nonspecific control (siC) (n=3 in duplicate). **(I)** Representative images show ARL13B and PCNT immunohistochemistry on day 2 of spontaneous differentiation in 3S-iPSCs transfected with the indicated siRNAs. **(J)** Si-RNA depletion of MCU in in 3S-iPSCs increases the fraction of cells with primary cilia (n = 3, 10 −16 fields per condition) (Scale bar = 10μm). **(K)** MCU knockdown in 3S-iPSCs increases PAX6+ medial fluorescence intensity on day 5 of differentiation (n=3, in duplicate). *Transcript levels are normalized to HPRT. Comparisons by ANOVA or Two-tailed T-test. Error bars represent SEM*. *\* P≤0.05, ** P ≤0.01, *** P ≤0.001, **** P ≤0.0001*

### Reduced commitment of DS 3S-iPSCs to NE is independent of sex and genetic background

The pair of isogenic lines used in our analysis were derived from the Coriell repository skin fibroblast cell line, AG05397, sampled from a 1-year-old male with DS and obtained through the NHLBI Progenitor Cell Biology Consortium (Parra *et al*., 2018). To validate our findings in an unrelated genetic background, we analyzed a pair of isogenic iPSCs derived from a 46-year-old female with DS (**Fig. EV5A**) that was generated by the Espinosa Laboratory at the Linda Crnic Institute for Down Syndrome (Klein *et al*, 2021). This pair of isogenic lines are designated here as 2SF-iPSC and 3SF-iPSC to distinguish them from the male lines. Consistent with the results from the male-derived lines, *ΔΨm* was higher in the 3SF-iPSCs than in the 2FS-iPSCs (**Fig. 5A**). The percent of 3SF-iPSCs in S phase was higher (p<0.0001), and the percent in G_1_/G_0_ was lower (p=0.0001) (**Fig. 5B**). Following 2 days of spontaneous differentiation, the fraction of cells with primary cilia was lower in the 3SF-derived cultures than in the euploid, 2SF-derived isogenic controls (**Fig. 5C and 5D**). After 5 days of differentiation, both *PAX6* transcript levels (**Fig. 5E)** and PAX6+ median fluorescence intensity **(Fig. 5F and 5G)** were lower in the 3SF-derived cultures. Pretreating the 3SF-iPSC with either the cell cycle inhibitor PD (**Fig. EV5B-EV5D**) or the MCU inhibitor Ru265 (**Fig. EV5B-EV5G**) increased the percentage of cells in G_1_/G_0_, as well as *PAX6* transcript levels and PAX6 median fluorescence intensity on day 5 of differentiation. Taken together, the female trisomic cells display the same differential properties observed in male 3S-iPSCs: those of increased mitochondrial activity, shorter cell cycle, reduced ciliation, and reduced commitment to NE when compared to their isogenic, euploid controls. Thus, the reduced capacity of DS iPSCs for early commitment toward the NE lineage appears to be independent of sex or genetic background, although this will be important to confirm this in additional lines, particularly those from DS individuals with partial trisomies.

**Figure 5:**
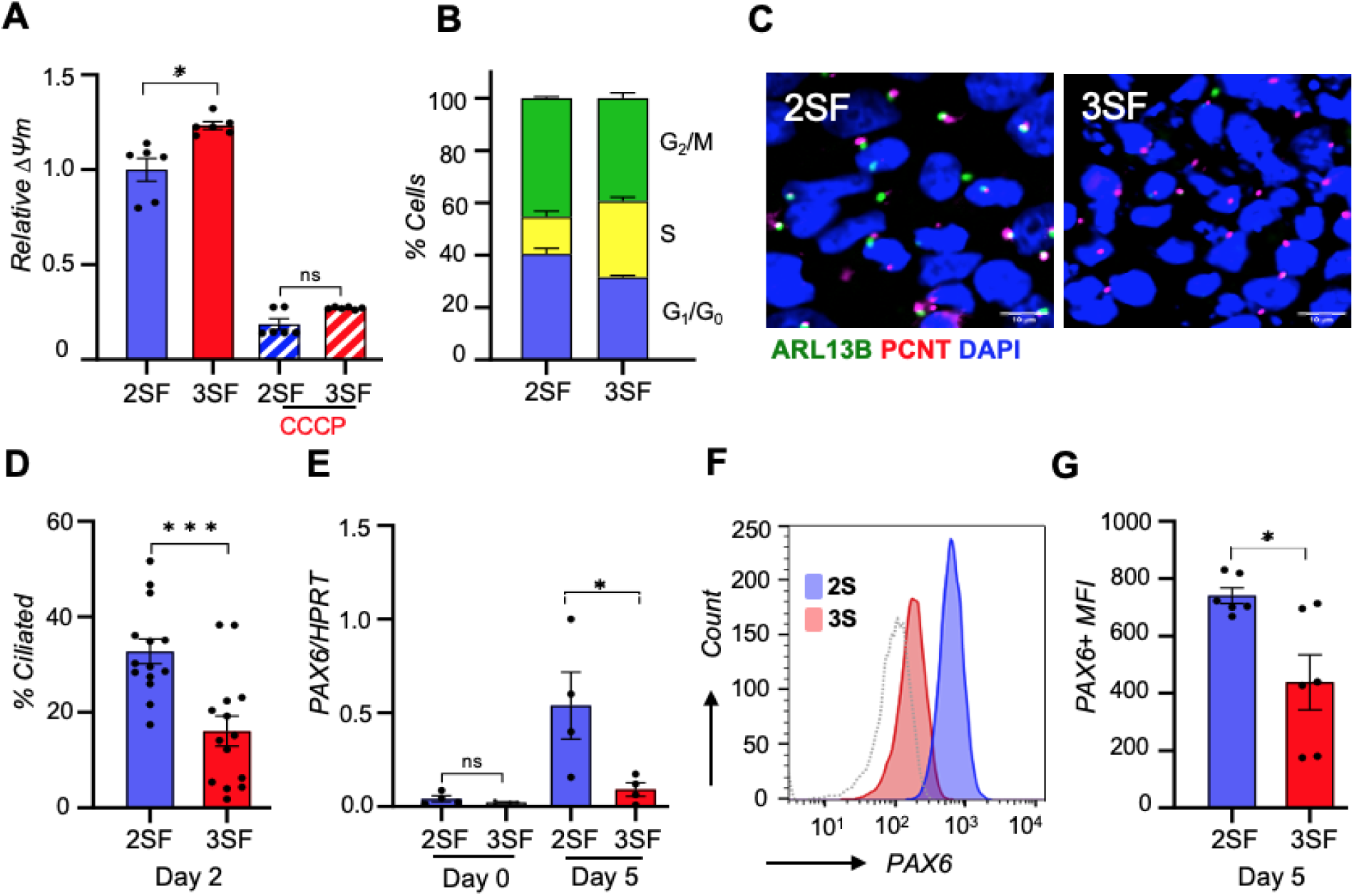
Ciliation and commitment to NE are also reduced in isogenic, female-derived, Down syndrome iPSCs. Key experiments are repeated here in isogenic, iPSCs derived from a female with DS, designated as 2SF-iPSC and 3SF-iPSC. **(A)** Mitochondrial membrane potential (*ΔΨm*) is higher in the 3SF-iPSCs than in 2SF-iPSCs. The addition of CCCP (20μM) is used as a negative control. (n=3. In duplicate) **(B)** The fraction of cells in G_1_/G_0_ is lower and the fraction in S phase higher in 3SF-iPSCs when compared to 2SF-iPSCs. **(C)** Representative images comparing ARL13B and PCTN staining of 2SF and 3SF cultures after 2 days of spontaneous differentiation. (Scale bar = 10μm). **(D)** Primary cilia are less abundant in the 3SF cultures than in the 2SF cultures on day 2 of spontaneous differentiation (n=12). **(E)** *PAX6* transcript levels and **(F and G)** PAX6+ median fluorescence intensity are lower in 3SF cultures than in 2SF cultures on day 5 of spontaneous differentiation (n=3, in duplicate). *Transcript levels are normalized to HPRT. Comparisons by ANOVA or Two-tailed T-test. Error bars represent SEM*. *\* P≤0.05, *** P ≤0.001*

### Pharmacological intervention can increase the commitment of DS 3S-iPSCs to NE

We tested a select group of small molecules for their ability to increase commitment of 3S-iPSCs to NE. Although structurally different, the three compounds we chose, fluoxetine, metformin, and nobiletin, have in common four key features: (1) They’ve been reported to increase neurogenesis and/or protect neurons from damage (Clark *et al*, 2006; Pang *et al*, 2023; Ruddy *et al*, 2019; Umemori *et al*, 2018; Wang *et al*, 2012; Xiong *et al*, 2023); (2) They’ve been reported capable of inhibiting mitochondrial activity, specifically electron transport complex I (ETCI) (Cikankova *et al*, 2020; Fontaine, 2018; Sharikadze *et al*, 2016); (3) They’ve been reported to promote cell cycle arrest at G_1_/G_0_ (He *et al*, 2024; Mogavero *et al*, 2017; Morley *et al*, 2007); and (4) they are widely used and well tolerated. Also, we found no reports of them impacting primary cilium form or function, as we wanted to void compounds known to disrupt proper function of this critical developmental signaling hub. We first tested fluoxetine, a serotonin selective reuptake inhibitor (SSRI), that has previously been shown to increase neurogenesis in the Ts65Dn mouse model of DS (Begenisic *et al*, 2014; Bianchi *et al*, 2010; Clark *et al*., 2006; Guidi *et al*, 2014). Pretreating 3S-iPSCs with 1 μM fluoxetine for 16 hours significantly increased the fraction of cells in G_1_/G_0_ compared to vehicle treatment *(p=0.004)* (**Fig. 6A**). The drug was then removed and cultures allowed to undergo spontaneous differentiation. Fluoxetine pretreatment significantly increased ciliation on day 2 of differentiation (**Fig. 6B and 6C**) as well as *PAX6* transcript levels **(Fig. S6B)** and PAX6+ median fluorescence intensity on day 5 of differentiation compared to vehicle treatment (**Fig. 6D).** Pretreatment with 10 mM metformin, a drug used to treat Type 2 diabetes, yielded similar results (**Figs. 6E-6H and Fig. EV6C**) as did pretreatment with 25μM of the plant flavonoid nobiletin (**Figs. 6I-6L and EV6D**). All three drugs also increased the fraction of cells in G_1_/G_0_ and commitment to NE in the female 3SF-iPSCs **(Fig. EV7)**. As was the case for PD and MCU inhibition, each of the three drugs also increased PAX6 levels in the 2S iPSC-derived euploid cultures **(Figs. 6D, 6H, 6K and EV6B-D)**. Unlike the response to PD and MCU inhibition, which increased PAX6 to comparable levels in both the 2S and 3S lines, the relative response of the euploid and trisomic cells to the three compounds was more variable. Pretreatment with metformin had a more pronounced impact on 3S-iSPCs than on 2S-iPSCs, whereas the pro-NE impact of fluoxetine and nobiletin was less prominent in the 3S line. In response to each of these drugs the response of the 3S lines was remarkably comparable to the 2S line treated with vehicle control. We checked the gating parameters from the flow analysis of PAX6+ signal and found evidence of a small, but significant increase in the percentage of low-FSC/low-SSC events in both the 2S and 3S cultures treated with metformin indicative of an increase in cell death (**Fig. EV8)**, eliminating metformin from further consideration.

**Figure 6:**
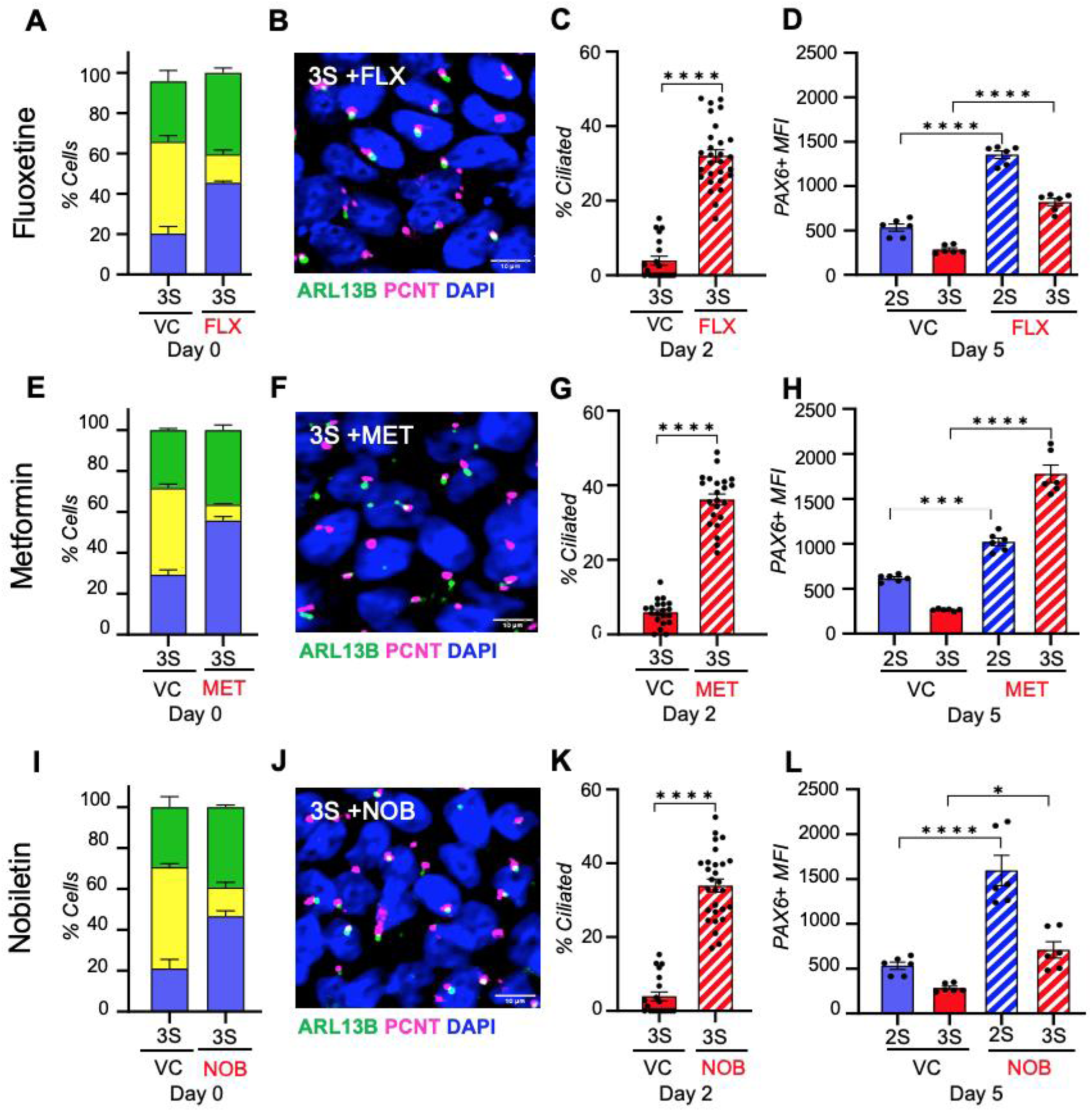
Pretreating 3S-iPSCs with small molecules increases NE commitment. **(A)** Pretreating 16-hours with 1μM fluoxetine (FLX) increases the fraction of 3S-iPSCs in G_1_/G_0_ compared to vehicle-treated controls (VC) (n=3, in duplicate). **(B)** Representative image of 3S-iPSCs pretreated with FLX shows an abundance of primary cilia on day 2 of spontaneous differentiation (Scale bar = 10μm). **(C)** FLX pretreatment increases the percentage of 3S cells with primary cilium. **(D)** FLX pretreatment increases PAX6+ median fluorescence intensity of both 2S and 3S derived cultures on day 5 of spontaneous differentiation (n=3, in duplicate). **(E)** Pretreating 16-hours with 10mM metformin (MET) increases the fraction of 3S-iPSCs in G_1_/G_0_ compared to vehicle-treated controls (VC) (n=3, in duplicate). **(F)** Representative image of 3S-iPSCs pretreated with MET shows an abundance of primary cilia on day 2 of spontaneous differentiation (Scale bar = 10μm). **(G)** MET pretreatment increases the percentage of 3S cells with primary cilium. **(H)** MET pretreatment increases PAX6+ median fluorescence intensity of both 2S and 3S derived cultures on day 5 of spontaneous differentiation (n=3, in duplicate). **(I)** Pretreating 16-hours with 25μM nobiletin (NOB) increases the fraction of 3S-iPSCs in G_1_/G_0_ compared to vehicle-treated controls (VC) (n=3, in duplicate). **(J)** Representative image of 3S-iPSCs pretreated with NOB shows an abundance of primary cilia on day 2 of spontaneous differentiation (Scale bar = 10μm). **(K)** NOB pretreatment increases the percentage of 3S cells with primary cilium. **(L)** FLX pretreatment increases PAX6+ median fluorescence intensity of both 2S and 3S derived cultures on day 5 of spontaneous differentiation (n=3, in duplicate). *Comparisons by ANOVA or Two-tailed T-test. Error bars represent SEM*. *\* P≤0.05, *** P ≤0.001, **** P ≤0.0001*.

Based on the data presented here and our previous analysis of mitochondrial structure and function in DS iPSCs (Parra *et al*., 2018), we propose the following model to describe a mitochondria-to-cilia signaling axis that functions early in lineage commitment and that is dysregulated in DS iPSCs (**Fig. 7**). Mitochondrial fission predominates in normal, euploid iPSCs (Parra *et al*., 2018), which are glycolytic with low metabolic activity, *ΔΨm,* and MCU capacity. Early in differentiation, cells begin to withdraw from cell cycle, increasing time spent in G_1_ and facilitating growth of a primary cilium that acts as a coordinating hub for signals directing lineage commitment to NE (Jang *et al*., 2016). Although DS iPSCs are fully capable of becoming NE, their trajectory is delayed due to elevated mitochondrial metabolic activity and MCU capacity that increases their rate of proliferation, thereby reducing time spent in G_1_ and growth of primary cilia required for commit to NE. Transient, early pharmacological interventions that slow the rate of proliferation and increase time spent in G_1_ are sufficient to increase commitment of DS iPSCs to NE *in vitro*.

**Figure 7:**
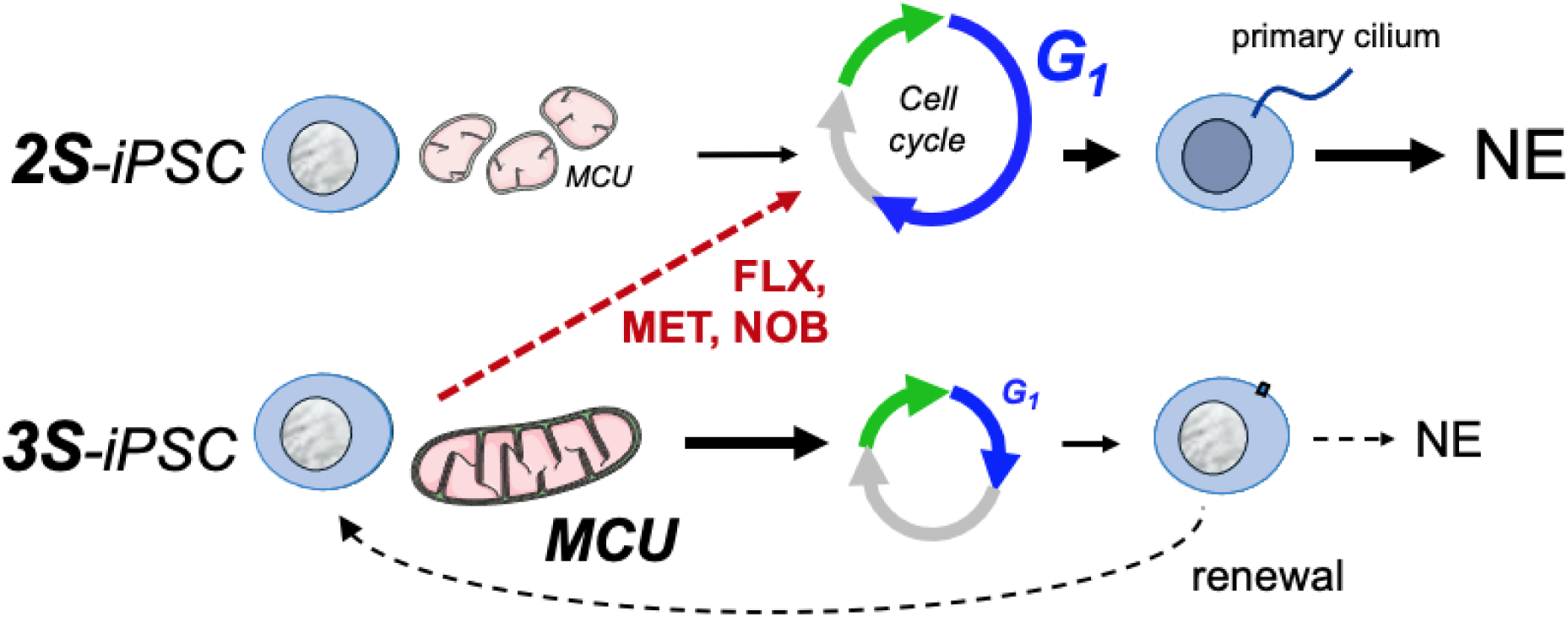
Model: In normal, euploid stem cells (2S-iPSC) the mitochondrial network is disrupted with low metabolic activity, ΔΨm, and MCU capacity. Early in lineage commitment, the cell cycle lengthens thereby increasing time spent in G_1_. This allows for growth of a primary cilium to act as a coordinating hub for signals directing commitment toward neuroectoderm (NE) and subsequent neurogenesis. The mitochondrial network in Down syndrome iPSCs (3S-iPSC) is more fused with higher oxidative metabolism, ΔΨm, and MCU capacity. Consequently, their rate of proliferation is more rapid with less time spent in G_1_, reducing the opportunity for growth of primary cilia and commitment to NE in favor of stem cell renewal. Pharmacological intervention with fluoxetine (FLX), Metformin (MET), or Nobiletin (NOB) slows proliferation, increasing the length of time spent in G_1_, and allowing growth of the primary cilium to improve commitment to NE.

## DISCUSSION

Cognitive disability is a universal feature of DS, with both reduced neurogenesis and changes in neuronal function contributing to deficits. Our studies demonstrate that iPSCs derived from individuals with DS have a reduced capacity to commit to NE, suggesting that a cell-autonomous property negatively impacts neurogenesis in DS at the very earliest stages of development.

During development of the mammalian central nervous system (CNS) gastrulation establishes the three primary germ cell layers: endoderm, mesoderm, and ectoderm. NE is derived from ectoderm and gives rise to neuroepithelial stem cells (NSCs), which ultimately give rise to all components of the CNS (Johnson *et al*, 2017). NSCs undergo symmetrical divisions to expand their own population then gradually transition to radial glial progenitor cells (RGCs), which initiate neurogenesis. RGCs initiate asymmetric divisions which both renew the pool of RGCs and generate either a neural-restricted progenitor cell (NPC) or a glial-restricted progenitor cell, primarily astrocyte progenitor cells (APCs) (**Fig. EV9)**. Ultimately it is the balance between expansion, self-renewal and consumption of the stem cell pools that establishes brain size. The production of new neurons ceases in most regions of the brain before, or shortly after birth, however, specialized niches of adult RGCs are established in the ventricular-subventricular zone and dentate gyrus of the hippocampus. These pools remain quiescent during development but can be reactivated for neurogenesis in the adult. Our studies examined commitment of iPSCs to NE, modeling an early event significantly upstream of commitment to either an NPC or APC populations. The exact relationship between the developmental trajectories of iPSCs *in vitro* and human development remains an ongoing debate, however, our findings are stikingly consistent with the well-established importance of cell cycle,(Salomoni & Calegari, 2010) mitochondrial dynamics,(Khacho & Slack, 2018) and primary cilia (Hasenpusch-Theil & Theil, 2021) in cell fate decisions later during neurogenesis and cortical development.

The length of the G1 phase of the cell cycle plays a central role in influencing the switch from expansion to differentiation of neural stem and progenitor cells (Salomoni & Calegari, 2010). There is an extensive literature demonstrating that antiproliferative genes promote neurogenesis, whereas positive regulators of proliferation have the opposite effect (Salomoni & Calegari, 2010). Our finding that more rapidly proliferating 3S-iPSCs are delayed in their commitment to NE is consistent with this as well as with *in vitro* studies showing that when hESCs are released to differentiate spontaneously, the cells within the population that commit to NE are those with longer cell cycles and an extended G_1_(Jang *et al*., 2016). Theoretically, a reduction or delay in commitment of DS stem cells to NE would be predicted to reduce or delay production of NSCs. How this might impact later stages of neurogenesis and whether it occurs *in vivo* will require further investigation.

Although our findings indicate that 3S-PSCs proliferated more rapidly than normal, this does not mean that this property translates directly to all subsequent lineages. NPCs(Murray *et al*., 2015; Tang *et al*., 2021) and APCs(Kawatani *et al*., 2021) derived from DS iPSCs are reported to have decreased and increased rates of proliferation respectively, yet they both derive from a common pool of NSCs. Interestingly, the observed decrease in proliferative potential of DS NPCs appears to require time to manifest as the percentage of neurons in neurospheres derived from euploid and trisomic iPSCs has been shown to be the same after 6 weeks of expansion, but lower in trisomic neurospheres after 10 weeks of expansion (Bhattacharyya *et al*, 2009). although not addressed in this paper, we postulate that the increased mitochondrial activity and associated increases in ROS could play a role in this later degenerative trajectory.

Our studies examine only the very early step of commitment to NE, however, here is emerging evidence that mitochondrial dynamics also play an active role in cell fate decisions of RGCs during both embryonic and adult neurogenesis (Khacho & Slack, 2018). Elegant studies have shown that following asymmetric the division of RGCs in the cerebral cortex, mitochondrial fission increases in the daughter cell destined to become a neuron, whereas mitochondrial fusion and oxidative metabolism increase in the daughter cell that retains stemness and returns to renew the pool of RGCs (Iwata *et al*, 2020). The Roede laboratory has shown that NPCs derived from DS iPSCs rely heavily on oxidative phosphorylation whereas normal, euploid NPCs remain primarily glycolytic (Prutton *et al*, 2023), similar to our observations of the metabolic characteristics of 3S-iPSCs prior to differentiation.

Metabolic and mitochondrial dysfunction are prominent themes in the scientific literature on DS. Some investigators report significant mitochondrial dysfunction (Mollo *et al*, 2020), whereas others report minimal dysfunction (Anderson *et al*, 2021). It is important to remember that mitochondrial dysfunction occurs whenever there is a mismatch between the production of ATP and energy demands in cells. Thus, over-active mitochondria can be as problematic as mitochondria generating insufficient ATP, and the observed metabolic phenotype may depend on the specific tissue and developmental time point one examines. For instance, hyperactivity of mitochondria in some neurons decreases neurotransmission and has a negative impact on behavior (Kanellopoulos *et al*, 2020). Hyperactive mitochondria have also been linked to Parkinson’s disease with suggestions that therapeutic intervention to reduce mitochondrial activity might be a new avenue for PD treatment (Mor & Murphy, 2020). Here we provide evidence that early in development elevated mitochondrial metabolism may delay neuronal development, however, one could also envision that over the course of a lifetime, increased ROS-generation, due to mitochondrial super-sufficiency, might accelerate age-related mitochondrial damage thereby contributing to degenerative aspects of DS. Consequently, interventions to reduce mitochondrial activity might be beneficial at some stages of development whereas improving mitochondrial function may be beneficial at others.

In our studies reduced ciliation in 3S-iPSCs correlated with a reduced ability to commitment to NE and each intervention that increased commitment to NE also restored ciliation. Primary cilia are present on RGCs in both the developing and adult cortex, where they influence multiple aspects of progenitor function(Hasenpusch-Theil & Theil, 2021; Zaidi *et al*, 2022). Primary cilia are also involved in the development of many other cell types and organ systems, including the heart. Congenital heart defects (CHDs) occur in approximately half of babies born with DS. Gene association studies in non-DS populations indicate that CHD-causing mutations frequently occur in genes related to the primary cilium (Gabriel *et al*, 2024). Using directed differentiation, we tested the ability of 3S-iPSCs to commitment to either endoderm or mesoderm, both of which contribute to specification and differentiation of the myocardium (Lough & Sugi, 2000). Commitment to either lineage was reduced when compared to the response of isogenic, 2S-iPSC controls (**Fig. EV10 and EV11**), raising the possibility that a generalized delay in the commitment of stem cells to any cell lineage could be a contributing factor in DS-associated CDH and other developmental delays. It may be relevant noting that a study comparing mitochondrial properties in normal fibroblasts to fibroblasts from DS patients reported elevated intra-mitochondrial Ca^2+^ and ROS production in the DS lines. Furthermore, the increase in mitochondrial Ca^2+^ and ROS were more pronounced in lines derived from DS individuals with CHD than in those derived DS individuals without CHD (Piccoli *et al*, 2013).

There are several cilia-related genes on HSA21 (Galati *et al*, 2018; Kogiso *et al*, 2020; Raveau *et al*., 2017), including, Pericentrin (PCNT), which has been shown to have a dose-dependent impact on the primary cilium (Jewett *et al*., 2023) and therefore increased dosage of PCNT likely impacts aspects of ciliation in the 3S-iPSCs. However, genetic analysis of DS patients with partial trisomy 21 suggests that triplication of PCNT is not required to manifest the intellectual disabilities and craniofacial features of DS (Pelleri *et al*, 2019). Furthermore, our experiments suggest that elevated mitochondrial metabolic activity lies upstream of reduced ciliation in 3S-iPSCs. Whether a specific gene, a combination of genes, or the state of aneuploidy itself is responsible for the increased mitochondrial metabolic activity of 3S-iPSCs and the reduction in ciliation remains to be determined. Importantly, our findings highlight mitochondrial metabolism and MCU capacity as a potential nodal point to target therapeutically, regardless of the specific genetic cause.

The three drugs we showed capable of increasing commitment of 3S-iPSCs to NE were chosen for testing in part because, although their primary targets are diverse, each has been reported to inhibit mitochondrial metabolism (Cikankova *et al*., 2020; Fontaine, 2018; Sharikadze *et al*., 2016). Although we show that pharmacological intervention can increase commitment to NE *in vitro,* our studies provide no insight into what an appropriate *in vivo* level of stem cell commitment to NE might be or when it might be beneficial. Extreme caution is warranted, and further studies are essential. Metformin provides an excellent example of the potential challenges. Although it has been found to increase neurogenesis in other *in vivo* animal models (Dadwal *et al*, 2015; Ruddy *et al*., 2019; Wang *et al*., 2012), daily administration of metformin to pregnant rats has a negative impact on neocortical development of the pups (Oner *et al*, 2024).

An important strength of our study is that it was carried out using human iPSCs rather than in a mouse model of DS. Although mouse models have proved an invaluable tool for studying may aspects of DS, they may not fully recapitulate the earliest deficiencies in neuronal development. Although in “newly born” cortical neurons mitochondria are initially fragmented, the mitochondrial network amplifies and fuses as they mature. The rate at which this occurs is species-specific even in culture, with the mitochondrial population in newly-formed human neurons taking several months to mature, whereas mouse neurons accomplish this in a matter of weeks (Iwata *et al*, 2023). CROCCP2 was recently identified as a human-specific modifier that promotes the expansion of cortical progenitors by decreasing ciliogenesis (Van Heurck *et al*, 2023). It is, therefore, possible that the mitochondria-to-cilia signaling axis we’ve proposed here may influence this critical expansion of cortical progenitors and have an even bigger impact on neurogenesis in humans than in mice. These differences will need to be taken into consideration during preclinical testing of any DS therapy in rodent models, which remains the most viable option for such studies.

Our model predicts that the reduction, or delay, in commitment of DS stem cells to NE would reduce the abundance of subsequent pools of progenitor cells, however, if DS stem cells are returning to renew the pool of stem cells rather than committing to differentiation, there may remain a viable pool of stem cells waiting to be mobilized buy as yet unidentified interventions able to normalize the balance between lineage commitment and self-renewal. While this is theoretically intriguing, in real life it would be impossible to apply such a therapy early enough to impact a point at which stem cells were committing to NE, however, if the mitochondria-to-cilia signaling axis we have described here also influences the behavior of RGCs in the developing and adult brain, then it could potentially emerge as a viable therapeutic target.

## Methods

### iPSC lines, origin and maintenance

Male isogenic 2S and 3S hiPSC lines were originally derived from a Coriell repository skin fibroblast line, AG05397, sampled from a 1-year-old male with DS. They were obtained through the NHLBI Progenitor Cell Biology Consortium and are described in (Parra *et al*., 2018). The pair of female isogenic line (2SF-iPSC and 3SF-iPSC) were generated at the Linda Crnic Institute for Down Syndrome from renal epithelial cells obtained from a 46-year old female with Down syndrome through the Human Trisome Project (COMIRB # 15-2170) using six factor RNA-based reprograming as described (Klein *et al*., 2021). In Klein et al they are designated as ILD11#3/ILD1(2)-1. They are renamed here to simply following their genetic identity in the text. As per International Society for Stem Cell Research (ISSCR) standards for human stem cell use in research, we regularly checked our iPSCs lines for pluripotency markers (by Immunofluorescence or qPCR or flowcytometry), mycoplasma contamination and periodically sent for karyotyping (Band resolution: 400-475) at the WiCell Research Institute, University of Wisconsin–Madison to verify genome stability. A Master Cell Bank was frozen back at feeder-free passage numbers between 12 and 23 then verified by karyotyping. If passage numbers exceeded 52 (or before) a new vial of cells would be obtained from the Master Cell Bank.

Cells were grown on Cultrex^TM^ BME-coated plates in mTeSR^TM^ Plus medium and passaged (splitting ratio minimum 1:6 and maximum 1:10) without enzymes using 0.5mM EDTA in PBS to disengage cells from plates. A ROCK inhibitor (10µM Y-27632) was added to the mTeSR Plus culture media during for the first 24 hours after splitting. When the cells reached 60% confluence, they were either passaged or used for experiments. All the experiments were carried out on karyotyped cells from the Master Cell Bank that were then passaged at least 4 to 5 times before the start of an experiment. Most all experiments were carried out at passage numbers between 18 and 52.

### Directed differentiation

The STEMdiff™ Trilineage Differentiation Kit (**Table 1**) from StemCell Technologies was used for directed differentiation of iPSCs into all three germ layers, ectoderm, mesoderm and endoderm. Briefly, iPSCs were released from plates by incubating in TrypLE Express (Gibco) at 37°C for 7 to 10 minutes. of incubation at 37°C. Single cells were counted and seeded on Cultrex-coated plates or chamber slides at the recommended cell densities. At 60% confluence cells were shifted to a lineage-specific medium, changing media daily for 5 days then harvested for transcript analysis, flow cytometry or immunofluorescence confocal microscopy.

**Table 1:**
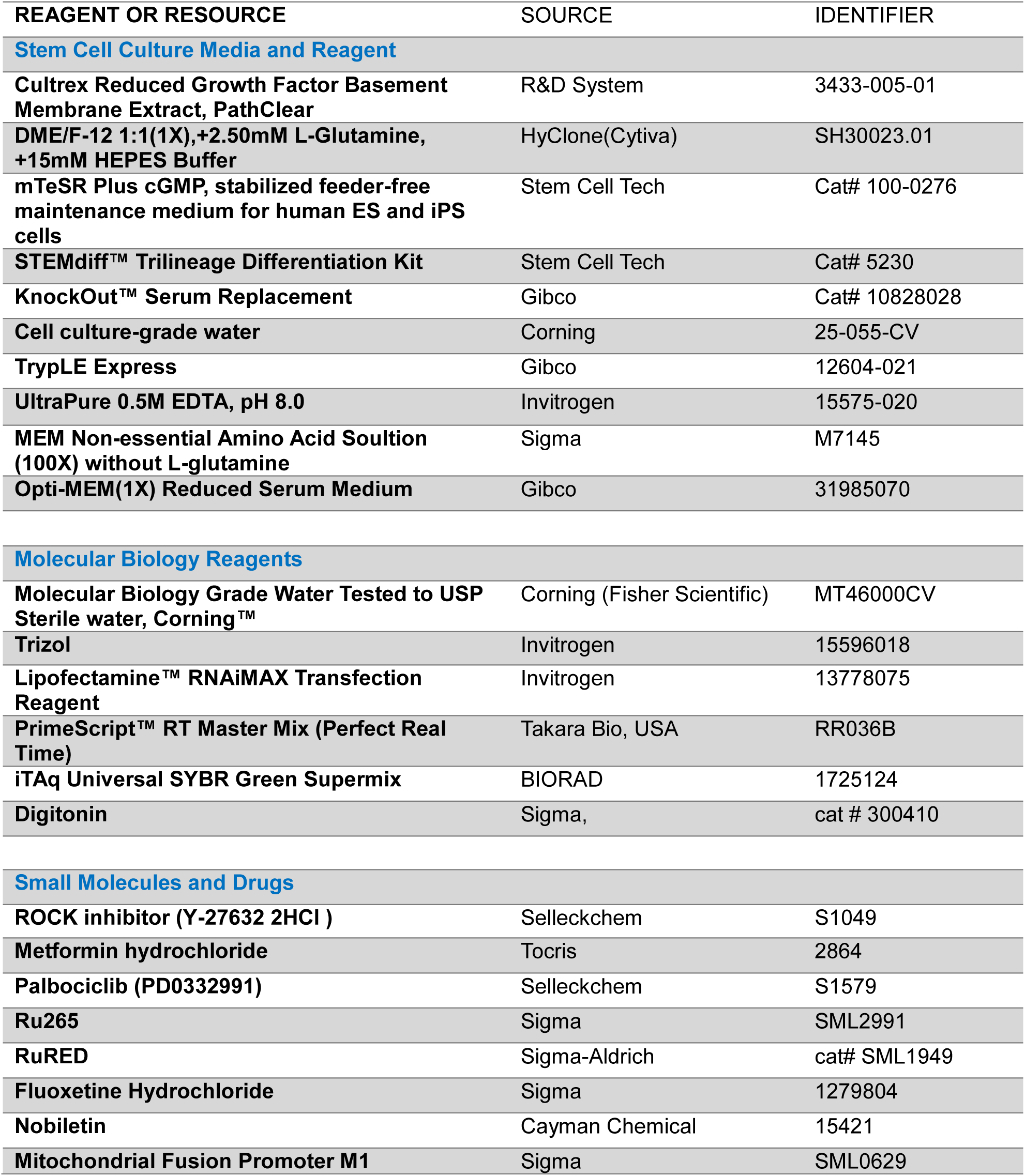
Details and sources of reagents.

| REAGENT OR RESOURCE | SOURCE | IDENTIFIER |
| --- | --- | --- |
| **Stem Cell Culture Media and Reagent** |  |  |
| Cultrex Reduced Growth Factor Basement Membrane Extract, PathClear | R&D System | 3433-005-01 |
| DME/F-12 1:1(1X),+2.50mM L-Glutamine, +15mM HEPES Buffer | HyClone(Cytiva) | SH30023.01 |
| mTeSR Plus cGMP, stabilized feeder-free maintenance medium for human ES and iPS cells | Stem Cell Tech | Cat# 100-0276 |
| STEMdiff™ Trilineage Differentiation Kit | Stem Cell Tech | Cat# 5230 |
| KnockOut™ Serum Replacement | Gibco | Cat# 10828028 |
| Cell culture-grade water | Corning | 25-055-CV |
| TrypLE Express | Gibco | 12604-021 |
| UltraPure 0.5M EDTA, pH 8.0 | Invitrogen | 15575-020 |
| MEM Non-essential Amino Acid Soultion (100X) without L-glutamine | Sigma | M7145 |
| Opti-MEM(1X) Reduced Serum Medium | Gibco | 31985070 |
| **Molecular Biology Reagents** |  |  |
| Molecular Biology Grade Water Tested to USP Sterile water, Corning™ | Corning (Fisher Scientific) | MT46000CV |
| Trizol | Invitrogen | 15596018 |
| Lipofectamine™ RNAiMAX Transfection Reagent | Invitrogen | 13778075 |
| PrimeScript™ RT Master Mix (Perfect Real Time) | Takara Bio, USA | RR036B |
| iTaq Universal SYBR Green Supermix | BIORAD | 1725124 |
| Digitonin | Sigma, | cat # 300410 |
| **Small Molecules and Drugs** |  |  |
| ROCK inhibitor (Y-27632 2HCl ) | Selleckchem | S1049 |
| Metformin hydrochloride | Tocris | 2864 |
| Palbociclib (PD0332991) | Selleckchem | S1579 |
| Ru265 | Sigma | SML2991 |
| RuRED | Sigma-Aldrich | cat# SML1949 |
| Fluoxetine Hydrochloride | Sigma | 1279804 |
| Nobiletin | Cayman Chemical | 15421 |
| Mitochondrial Fusion Promoter M1 | Sigma | SML0629 |

### Spontaneous Differentiation

To allow spontaneous differentiation, iPSCs were seeded (1:10 splitting ratio) on Cultrex-coated plates. At 40 to 50% confluency, mTesR^TM^ Plus was replaced with media composed of DMEM/F12, supplemented with 15% knockout serum replacement, 1X MEM non-essential amino acids, and 0.1 mM β-mercaptoethanol. Medium was replaced every other day. Pharmacological interventions or vehicle controls were added to cultures at 40% confluency, pretreating for 16 hours prior to changing to spontaneous differentiation media without drug. Drug interventions included 5μM Palbociclib, 10nM Ru265, 1μM Fluoxetine, 10mM Metformin, or 25μM Nobelitin.

### siRNA transfection

Lipofectamine™ RNAiMAX Transfection Reagent was used according to instructions. Briefly, cells were plated on Cultrex-coated plates 6-well plates at a 1:10 splitting ratio and allowed to reach 50-60% confluency. Prior to transfection, cells were pretreated for an hour with the addition of 10 µM Y27632. Cells were washed once with Optimem® then incubated at room temp with 10 nM (10 pmol/ml) of siRNA of interest in 400 μL of Optimem®. After 20 minutes mTESR^TM^ Plus was added to bring the culture volume to 1 ml, which was changed 24 hours later to fresh mTESR^TM^ Plus. Knockdown efficiency was determined by qRT-PCR of the respective target gene normalized to *HPRT*. 48 hours after siRNA transfection, were shifted to spontaneous differentiation media. Details of siRNAs are provided in **Table 2**. Two different siRNAs where used for each gene targeted, validated for KD separately and then used in 1:1 combination in experiments.

**Table 2:** siRNAs used in studies.

| siRNA ID | Gene Symbol | Species | Prediction model | Company |
| --- | --- | --- | --- | --- |
| NM_138357_SASI_Hs01_00234628 | *MCU* | Human | Rosetta Prediction | Sigma |
| NM_138357_SASI_Hs01_00234629 | *MCU* | Human | Rosetta Prediction | Sigma |
| NM_007054_SASI_Hs01_00208830 | *KIF3A* | Human | Rosetta Prediction | Sigma |
| NM_007054_SASI_Hs01_00208831 | *KIF3A* | Human | Rosetta Prediction | Sigma |
| NM_174887_SASI_Hs01_00186053 | *IFT20* | Human | Rosetta Prediction | Sigma |
| NM_174887_SASI_Hs01_00186054 | *IFT20* | Human | Rosetta Prediction | Sigma |
| MISSION® siRNA Fluorescent Universal Negative Control #2, 6-FAM (SIC008-10NMOL) | Negative control | Human |  | Sigma |

### Immunofluorescence

Cells were fixed with 2% paraformaldehyde (diluted in PBS w/o Ca++ and Mg++, pH 7.4) for 10mins followed by permeabilization with 0.5% Triton X-100 (in PBS, pH 7.4, without Ca^2+^ or Mg^2+^) for 10 minutes at room temperature. Cells were washed gently three times using 1X Cyto-Fast^TM^ Perm/wash solution (Biolegend, USA) then incubated in serum-free protein-blocking solution (Abcam, ab64226) for 30 minutes to prevent non-specific antibody binding. Following blocking, cells were washed once with PBS then incubated overnight at 4C in a humidified chamber with primary antibodies (1:200), as listed in **Table 3**. The following day cells were washed three times with PBS then incubated for 1 hr at room temperature with the appropriate fluorophore-conjugated secondary antibodies (1:300), as listed in **Table 3**. Cells were again washed three times with PBS and then mounted for imaging using ProLong™ Gold Antifade reagent with DAPI (Invitrogen) to stain DNA. Images were acquired using a Nikon A1R Confocal microscope with NIS-Element AR 3.22.11 software.

**Table 3:** Antibodies.

| REAGENT OR RESOURCE | SOURCE | Dilution | IDENTIFIER |
| --- | --- | --- | --- |
| **Antibodies** |  |  |  |
| **Anti-OCT4 (Oct3) Antibody** | Biolegend | 1:200 | Cat# 653701 |
| **Anti-SSEA-4 Antibody** | Biolegend | 1:500 | Cat# 330401 |
| **Anti-NANOG Antibody** | Biolegend | 1:200 | Cat# 674201 |
| **Anti-SOX2 Antibody-PE** | Biolegend | 1:200 | Cat# 656103 |
| **Anti-PAX6 Antibody** | Biolegend | 1:200 | Cat# 862001 |
| **Anti-SOX17 Antibody** | Biolegend | 1:200 | Cat# 698501 |
| **Anti GATA6 antibody** | Novus Biologicals | 1:200 | Cat# MAB1700 |
| **Anti MESP1 antibody** | Novus Biologicals | 1:200 | Cat# MAB9219 |
| **Anti-Pericentrin antibody** | Novus Biologicals | 1:200 | Cat# NBP2-97596 |
| **Anti-OTX2 Antibody** | R&D | 1:200 | Cat# AF1979 |
| **Anti ARL13B antibody** | Santa Cruz | 1:200 | Cat# SC-515784 |
| **Alexa Fluor® 647 anti-mouse IgG2b** | Biolegend | 1:300 | Cat# 406718 |
| **Alexa Fluor® 488 anti-mouse IgG2b** | Biolegend | 1:300 | Cat# 406634 |
| **Alexa Fluor® 647 anti-mouse IgG2a** | Biolegend | 1:300 | Cat# 407116 |
| **Alexa Fluor® 488 anti-mouse IgG2a** | Biolegend | 1:300 | Cat# 407122 |
| **Alexa Fluor® 594 anti-mouse IgG1** | Biolegend | 1:300 | Cat# 406507 |
| **Alexa Fluor® 488 anti-mouse IgG1** | Biolegend | 1:300 | Cat# 406626 |
| **Alexa Fluor® 647 Donkey anti-rabbit IgG** | Biolegend | 1:300 | Cat# 406414 |
| **Alexa Fluor® 488 Donkey anti-rabbit IgG** | Biolegend | 1:300 | Cat# 406416 |
| **Staining Reagents** |  |  |  |
| **DAPI ready-made solution** | Sigma |  | Cat# MBD0015-5ML |
| **Protein Block** | Abcam |  | Cat# ab64226 |
| **Cell Staining Buffer** | Biolegend |  | Cat# 420201 |
| **BD Cytofix/Cytoperm™ Fixation/Permeabilization Solution Kit** | BD |  | Cat# BDB554714 |
| **TMRM** | Invitrogen |  | T668 |
| **MitoSOX-RED** | Invitrogen |  | M36008 |
| **CCK8 Cell counting kit** | APEX BIO |  | K1018 |
| **ProLong Gold antifade reagent with DAPI** | Invitrogen |  | P36935 |
| **DPBS w/o Calcium Chloride and Magnesium chloride** | Sigma-Aldrich |  | D8537 |
| **Rhod-2-AM** | Thermo Fisher |  | R1245MP |

### Image Analysis and cilia quantification

FIJI software (Schindelin et al., 2012) was used for image analysis. To assess ciliation multiple images were obtained of DAPI-stained cells reacted with antibodies against Pericentrin (PCNT) to detect centrosomes and ARL13B to mark primary cilia. A blinded scoring strategy was used to count ARL13B/PCNT double-positive cells divided by the total number of DAPI-positive cells. More than 2,000 cells were counted per experimental condition.

### RNA isolation and quantitative real-time PCR

Total RNA was isolated using Trizol reagent (Invitrogen) according to the manufacturer’s protocol. Isolated RNA was reverse transcribed with the 5X PrimeScript RT MasterMix (TaKaRa). RT-qPCR was performed on a C1000 Thermal Cycler CFX96 (Bio-Rad) using iTaq Universal SYBR Green Supermix (Bio-Rad). See **Table 4** for primers used in this study. Transcript levels were normalized to *HPRT*.

**Table 4:**
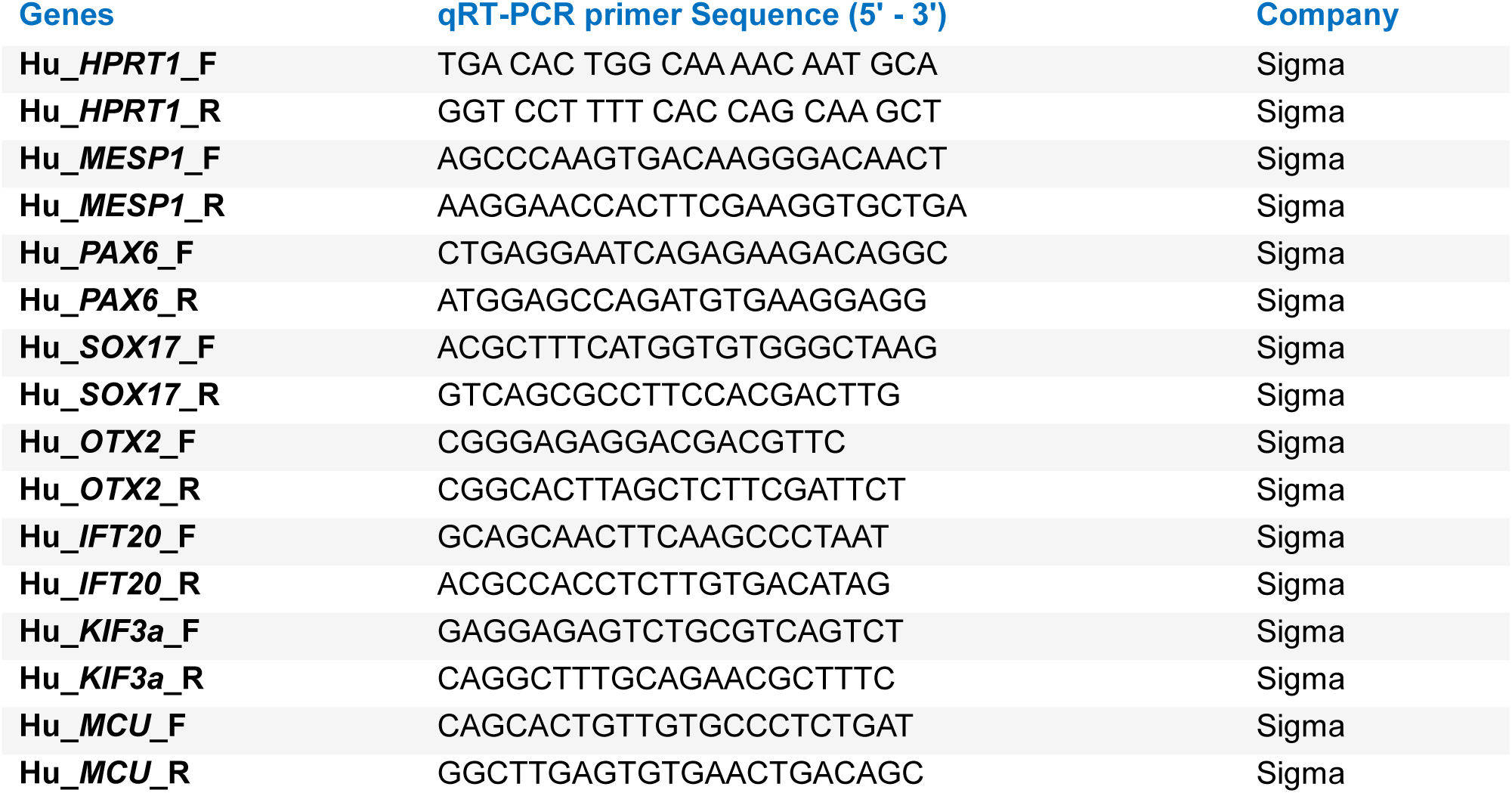
Primers.

### Cell Proliferation assay

Changes in cell density were quantified using the CCK8 Cell Counting Kit from APExBIO following the manufacturer’s instructions. Briefly, cells were seeded in 96-well plates at a concentration of 5,000 cells per well. A baseline reading was obtained on day then obtained daily for 3 consecutive days. Results were first normalized based on the 0^th^-day reading for that genotype and then normalized as a fold relative to the euploid reading at day 0.

### Cell cycle analysis by flow cytometry

One parameter flow cytometric analysis based on DNA content was used to determine the proportional distribution of cells across the cell cycle. Incubation of adherent cells with TrypLE Express for 7 to 10 minutes at 37°C was used to obtain single cells, which were then washed with PBS (pH 7.4, without Ca2+ or Mg2+), spinning at 1500 rpm in a benchtop IEC Centra CL2 Centrifuge for 5 minutes in a 15 ml conical tube. The cell pellets were fixed in 4 ml of ice-cold 75% ethanol vortexing vigorously and then stored at - 20°C for a minimum of 30 minutes or until the day of analysis. Prior to flow analysis, cells were pelleted again by centrifugation to remove ethanol, followed by two washes with PBS. They were resuspended in a PBS containing 20 μg/ml propidium iodide (PI) and 10–50 μg/ml RNase A then maintained at 4°C for a minimum of 30 minutes prior to flow. Fluorescence (FL2) was measured using a BD FACS Canto flow cytometer, and FlowJo 4.1 software used to determine the cell cycle phase distribution with the help of the manufacturer supplied Dean-Jett-Fox model.

### Flow cytometry analysis of linage markers

Cells were harvested with TrypLE Express, transferred to a 15ml conical tube, pelleted by centrifuging for 5 min at 1500 rpm then washed with 2ml of ice-cold PBS (pH 7.4, without Ca^2+^ or Mg^2+^) supplemented with 0.1% bovine serum albumin, and pelleted again. Ice-cold 75% methanol was used to fix and permeabilize the cells, which were maintained in 75% methanol for at least 30 minutes or stored at −20°C until ready for flow analysis. For antibody staining, samples were transferred to a 1.5 ml Eppendorf, centrifuged for 5 minutes at 4° C, 1500 rpm/212 rcf to remove the ethanol, and then washed twice with ice-cold PBS/0.1% BSA. Samples were incubated with primary antibody **(Table 3)** diluted 1:200 in 100 µL of PBS at room temperature for 1-2 hrs, then washed twice with PBS followed by incubation in the appropriate fluorochrome-conjugated secondary antibody (1:300 in PBS) for 1 hour at room temperature. Antibody was removed, samples were washed twice with PBS and resuspended in 600 µL of PBS. The BD FACS Canto flow cytometer was calibrated using UltraComp eBeads compensation beads (Invitrogen) prior to running samples. Results were analyzed using FlowJo Software v10.8.1.

### Flow analysis of mitochondrial membrane potential and ROS

Non-quenching levels of TMRM were used to assess mitochondrial membrane potential (*ΔΨm*). Live cells were loaded with 50 nM TMRM for 30 min; excitation/emission (543/560). Cells were trypsinized with TrypLE Express to remove them from the plate, washed twice with PBS, and fluorescence assessed by flow cytometry using a FACS Canto system (Becton Dickinson, Franklin Lakes, NJ, U.S.A). The addition of an uncoupler CCCP (20 μM) was used as a negative control. ROS levels were assessed similarly by loading live cells for 10 minutes with 5 μM MitoSox®; excitation/emission 510/580) and processing as described for TMRM.

### Assessment of MCU capacity

MCU capacity was assessed using the acetoxymethyl-ester Ca^2+^ probe, Rhod-2-AM (Thermo Fisher, cat #R1245MP) to track changes in mitochondrial Ca^2+^ levels ([Ca^2+^]mt) in permeabilized iPSCs as previously described (Maxwell *et al*, 2018; Parra *et al*., 2018). Briefly, iPSCs were incubated for 30 min at room temperature (20-22°C) with 2 µM Rhod-2-AM in a modified Tyrode Salt’s solution composed of: 140.0 mM NaCl; 3.7 mM KCl; 2.0 mM CaCl_2_; 1.0 mM MgCl_2_; 5.0 mM D-glucose; 10 mM HEPES and 0.5 mM NaH_2_PO_4_, pH 7.4 (NaOH). The “intracellular-like solution” mimicking cytosolic composition and osmolarity, was composed of: 120.0 mM KCl, 5.0 mM Sodium Succinate, 5.0 mM Sodium Pyruvate, 1.0 mM KH_2_PO_4_, 1.0 mM MgCl_2_; 20.0 mM HEPES, pH 7.4 (KOH). The intracellular-like solution was supplemented with either 1 mM EGTA (for Ca^2+^-free conditions) or 10 µM EGTA-buffered free-Ca^2+^ (calculated using MAXCHELATOR software (pH = 7.4, 22 C, 0.162 N, Ionic Contribution [ABS] 0.0020199 N) (Yoast *et al*, 2021). Cells were permeabilized by perfusing with 20 µM Digitonin in intracellular-like +EGTA solution for 30 seconds, followed by 30 seconds of perfusion with intracellular-like +EGTA solution without Digitonin. MCU was initiated by bringing external free-Ca^2+^ levels to 10 µM then recording Rhod-2-AM fluorescence for the next 100s. The identity of MCU was confirmed by the addition of 10 μM of the MCU inhibitor Ruthenium Red (RuRed) Ca^2+^ transients were measured at room temperature (20-22°C) using a polycarbonate 260 μL chamber (Harvard Bioscience Inc., USA cat #RC21BR) mounted on an inverted microscope coupled to a perfusion system electrically controlled. Fluorescence recordings were obtained using the Nikon Ti2-E Eclipse inverted light microscope, equipped with an objective (Nikon S Fluor × 20; numerical aperture: 0.75) and a digital SLR camera (DS-Qi2; Nikon, Japan) controlled by computer software (NIS Elements version 5.20.01, USA). iPSC cells were continuously perfused by a six-way perfusion system (VC-8 valve controller) at 4-5 ml per min with a common outlet 0.28-mm tube driven by controlled valves (Harvard Bioscience Inc., USA). Emitted fluorescence was collected through a 555 nm emission filter. All fluorescence images were generated at 5s intervals and normalized, and the values were calculated using Image J (1.53J).

### Statistics and reproducibility

Statistical analysis was performed using the GraphPad Prism software version 9.4.1 (GraphPad Software, San Diego, CA, USA) or Microsoft Excel (Microsoft® Excel® for Microsoft 365 MSO (Version 2408 Build 16.0.17928.20114) 64-bi). Analysis of variance (ANOVA) with Bonferroni and Šídák post hoc test and t-test were used to evaluate the statistical significance. P < 0.05 was defined to be significant. A minimum of three experiments on 3 different days were performed for each experimental set-up.

## RESOURCE AVAILABILITY

### Lead Contact

Further information and requests for resources and reagents should be directed to and will be fulfilled by the lead contact, Beverly Rothermel.

### Materials availability

This study did not generate new unique reagents.

### Data and code availability

Any additional information required to reanalyze the data reported ion this paper is available from the lead contact upon request.

## ACKNOWLEDGMENTS

This work was supported by NIH grants R01 HD101006 to B.A.R., 2R01NS055028 to W.L and B.A.R., R01DE027679 and R01DE032846 to R.S.L., and the Global Down Syndrome Foundation.

## AUTHOR CONTRIBUTIONS

Conceptualization, M.C. and B.A.R.; Methodology, M.C., B.A.R, and R.S.L.; Investigation, M.C, G.H.S.B., N.J., Y.D., and B.A.R.; Resources, B.F.N. and J.M.E.; Writing –Review &

Editing, M.C., B.A.R, B.F.N., J.M.E., E.R., W.L. R.S.L.; Funding Acquisition, B.A.R., R,S.L., W.L., and J.M.E.

---

**Figure EV1:**
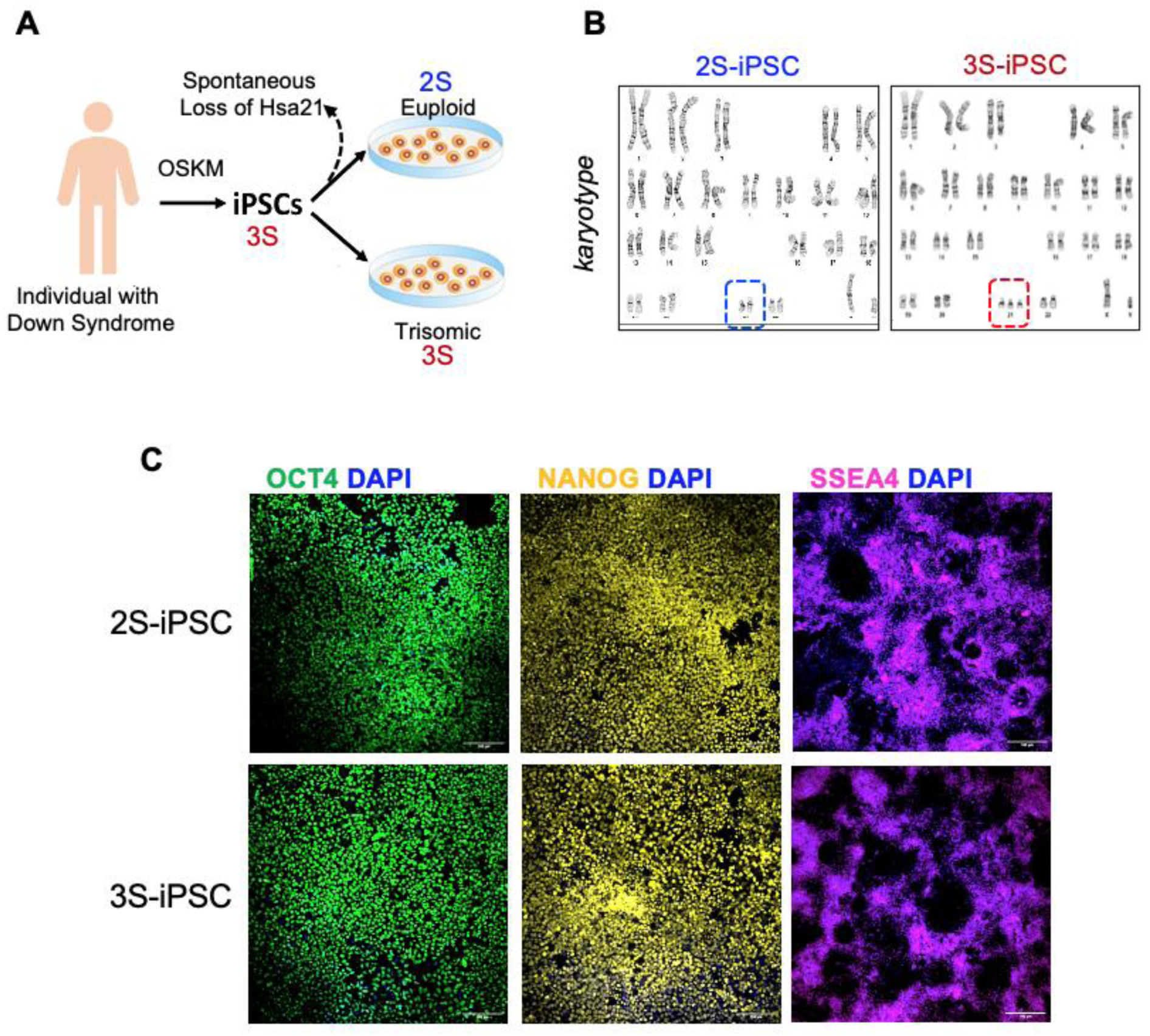
**(A) Schematic showing derivation of the isogenic iPSCs:** The isogenic, 2S/3S iPSC lines we obtained for use in our studies had been generated by OSKM (OCT4, SOX2, KLF4, MYC) reprograming of DS cells followed by passaging to isolate and expand clones that had lost the extra copy of HSA21, which occurs spontaneously at a low frequency. **(B)** Karyotyping was carried out periodically to verify genomic integrity and maintenance of the extra chromosome. (**C)** Immunohistochemistry for the pluripotency markers OCT4, NANOG and SSEA4 showed uniform staining in both the eupoid 2S and trisomic 3S iPSC lines. (Scale bar = 50 μm).

**Figure EV2:**
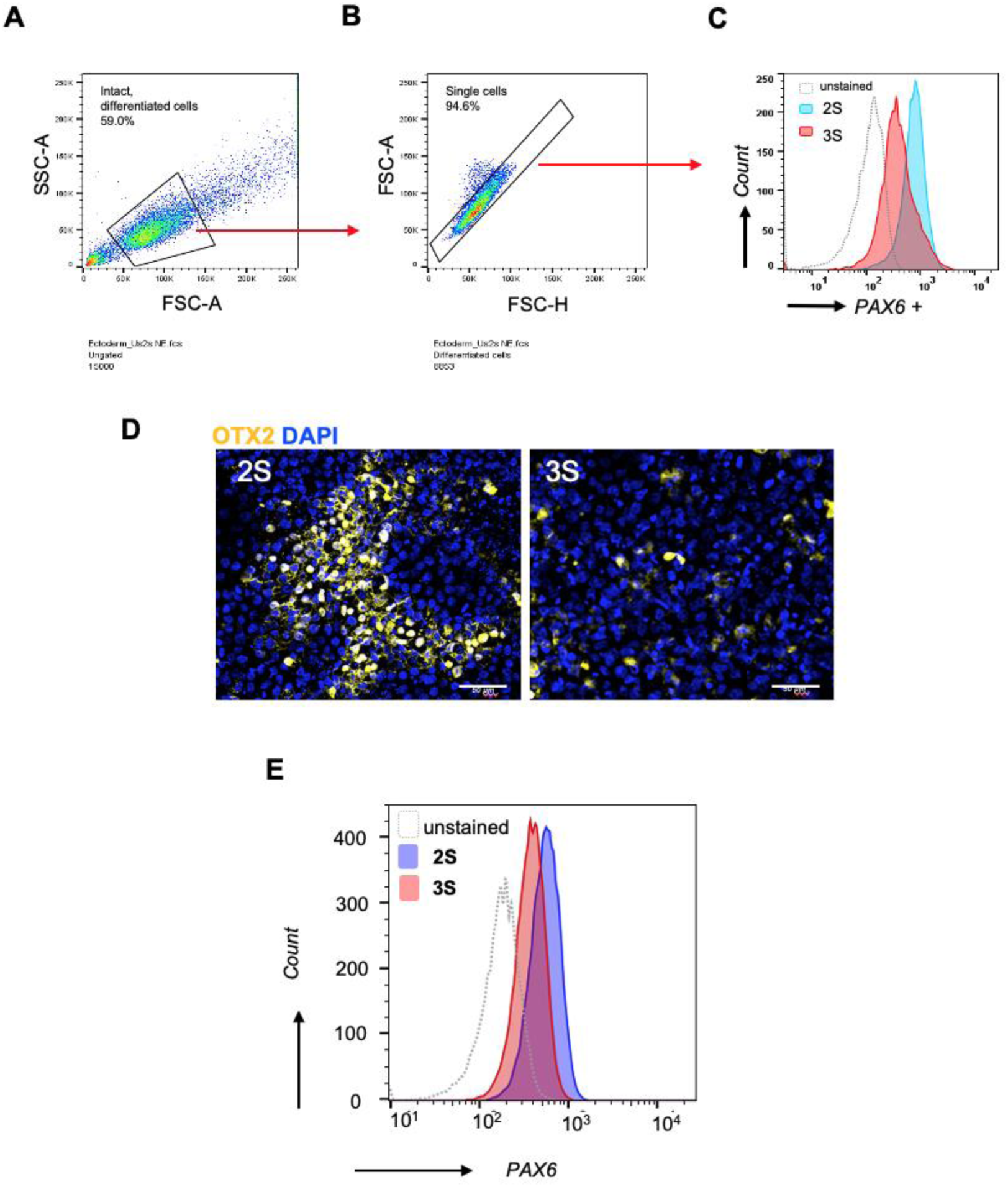
Flow cytometry gating strategy used to quantify PAX6+ signal: On day 5 of differentiation, single cells were dissociated from the plate and fixed, then stained with an antibody against PAX6 followed by a fluorescent-tagged secondary antibody. Pannels A and B show the gating strategies used to select the population of intact, differentiated cells **(A)** and to discriminate single cells from doublets **(B).** The histogram overlay in **(C)** compares the intensity of PAX6+ signal after 5 days of spontaneous differentiation. Unstained 2S-iPSCs not reacted with the primary antibody were used as a negative control. **(D)** 2S and 3S-iPSC cultures were allowed to undergo spontaneous differentiation for 5 days, then fixed and stained for OTX2 (yellow). DAPI-stained nuclei are in blue (Scale bar = 50μm). **(E)** Flow analysis of PAX6 stained cultures shows lower PAX6+ median fluorescence intensity in the 3S-derived cultures than in the 2S-derived cultures on day 5 of differentiation.

**Figure EV3:**
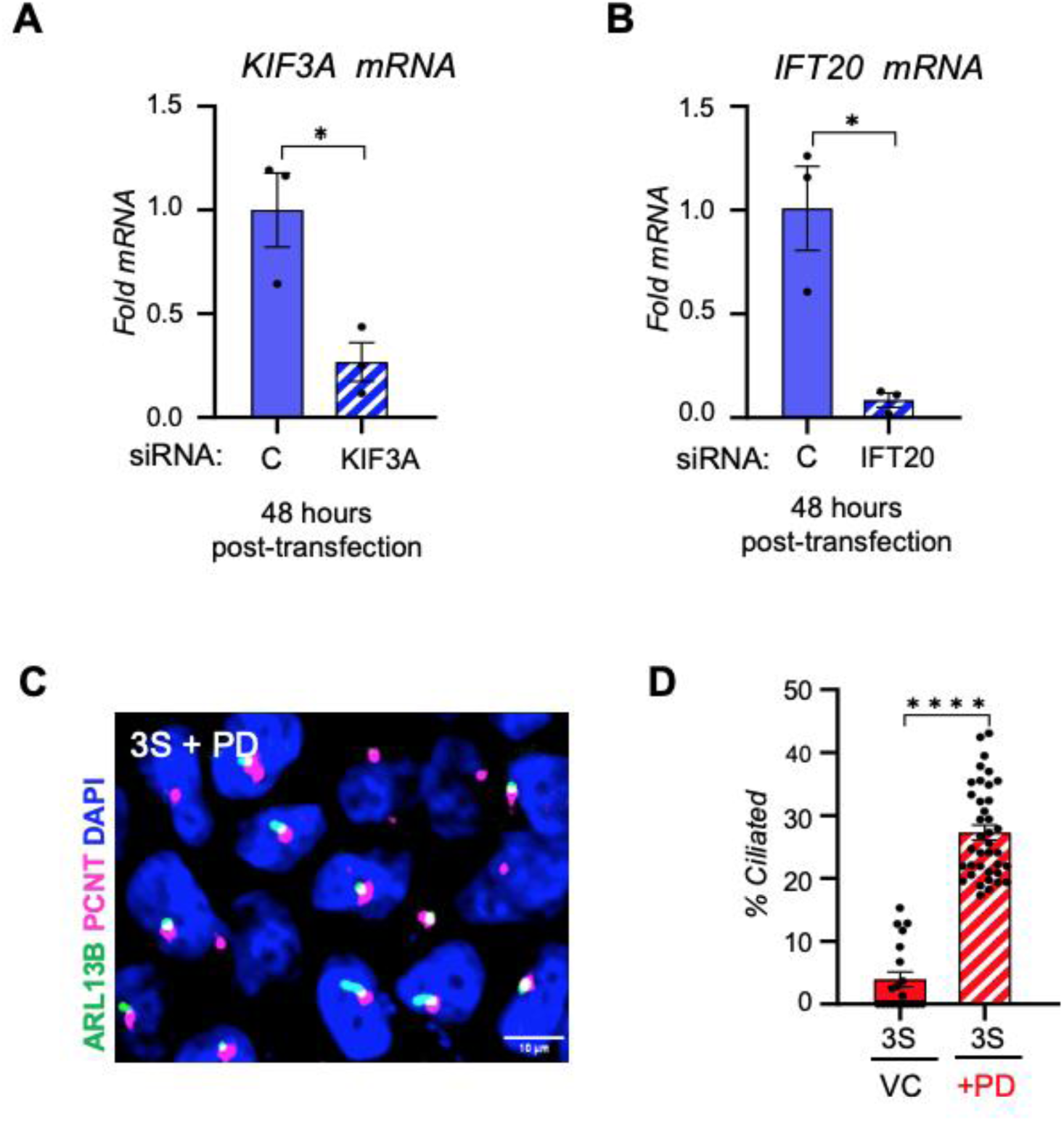
Si-RNA-mediated Knockdown of *KIF3A* and *IFT20* in 2S-iPSCs: Quantitative-RT-PCR was used to monitor siRNA-depletion of *KIF3A* **(A)** and *IFT20* **(B)** transcripts 48 hours after si-RNA transfection (n=3). **(C)** Representative image depicts ARL13B+ and PCTN+ staining of 3S-derived cultures that were pretreated for 16 hours with 5µM of the cell cycle inhibitor PD0332991 (PD) and then allowed to differentiate spontaneously for 2 days. (Scale bar = 50 μm). **(D)** Pretreatment of 3S-iPSCs with PD increased ciliation. (n=3 independent experiments, each dot represents quantification of a field of cells). *Comparisons by Two-tailed T-test. Error bars represent SEM*. *\* P≤0.05,,**** P ≤0.0001*

**Figure EV4:**
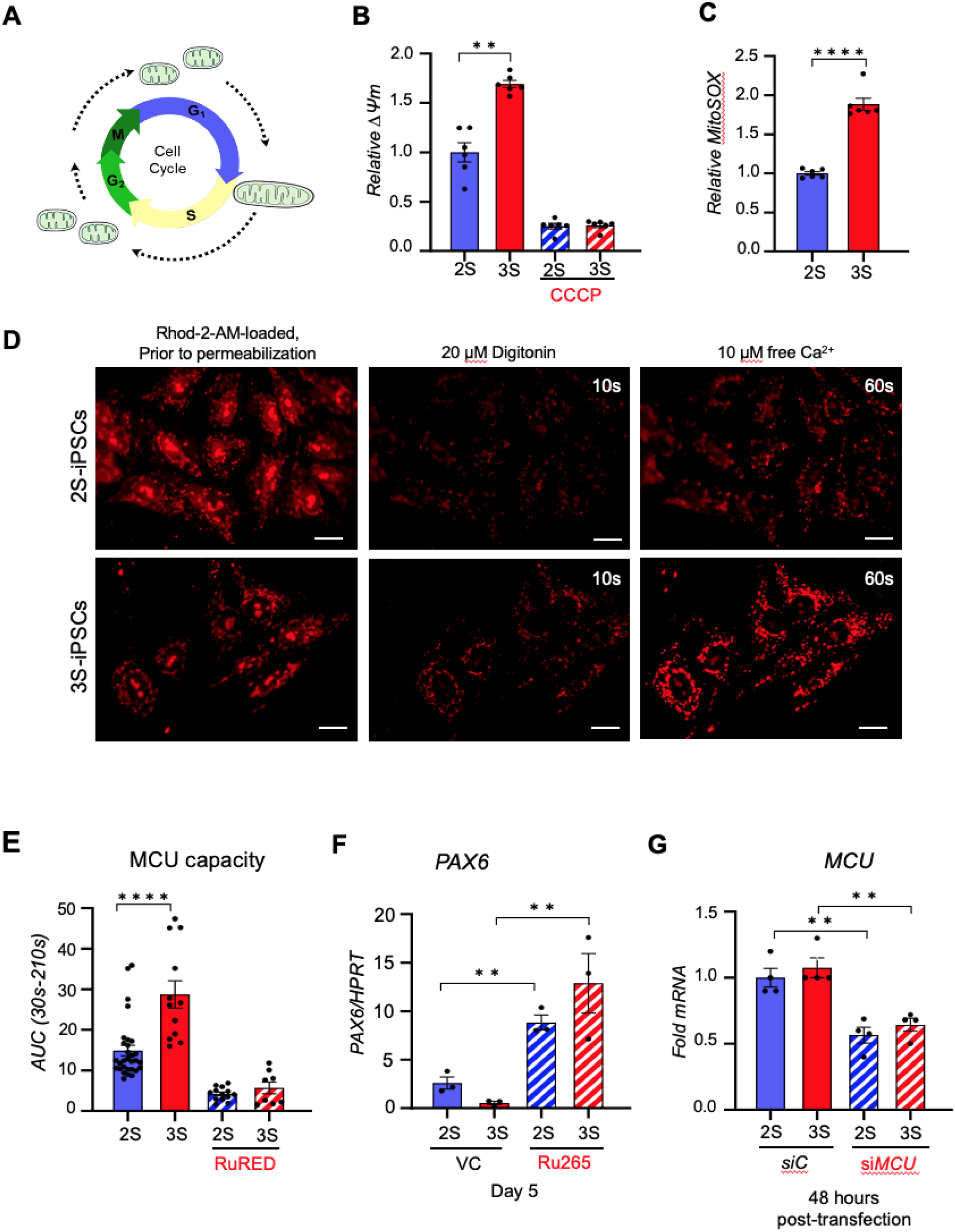
Studies related to MCU: **(A)** Schematic depicting mitochondrial fusion at the G_1_/S transition **(B)** Mitochondrial membrane potential (*ΔΨm*), measured by flow, is higher in 3S-iPSCs than in 2S-iPSCs, measured in live cells loaded with 50 nM TMRM. CCCP (20μM) was added as a negative control. (n=6). **(C)** MitoSOX-Red intensity is higher in 3S-iPSCs than in 2S-iPSCs (n=6). **(D)** Representative images of F_555_ signal in iPSCs loaded with Rhod-2-AM prior to permeabilization, 10 seconds after permeabilization, and 50 seconds after the addition of 10μM free Ca^2+^. **(E)** MCU capacity, quantified as the area under the curve (AUC) between 30 seconds and 210 seconds, was greater in 3S-iPSCs than in isogenic, 2S-iPSCs and abolished by the addition of 10μM RuRED (+RR) (n=3 independent experiments and dots represent data from individual cells. **(F)** *PAX6* transcript levels were higher in 2S and 3S-iPSCs pretreated for 16 hours with the MCU inhibitor Ru265 (10nM) on day 5 of spontaneous differentiation compared to vehicle-treated controls (VC) (n=3 independent experiments in duplicate). **(G)** siRNA-mediated targeting of MCU reduced transcript levels 48 hours after transfection. (n=4). Transcript levels are normalized to *HPRT.* In G *Error bars represent SEM*. Significance was determined by one-way ANOVA followed by Tukey’s multiple comparison post-hoc test, or a Two-Tailed T-test. *\* P≤0.05, ** P ≤0.01, *** P ≤0.001, **** P ≤0.0001*.

**Figure EV5:**
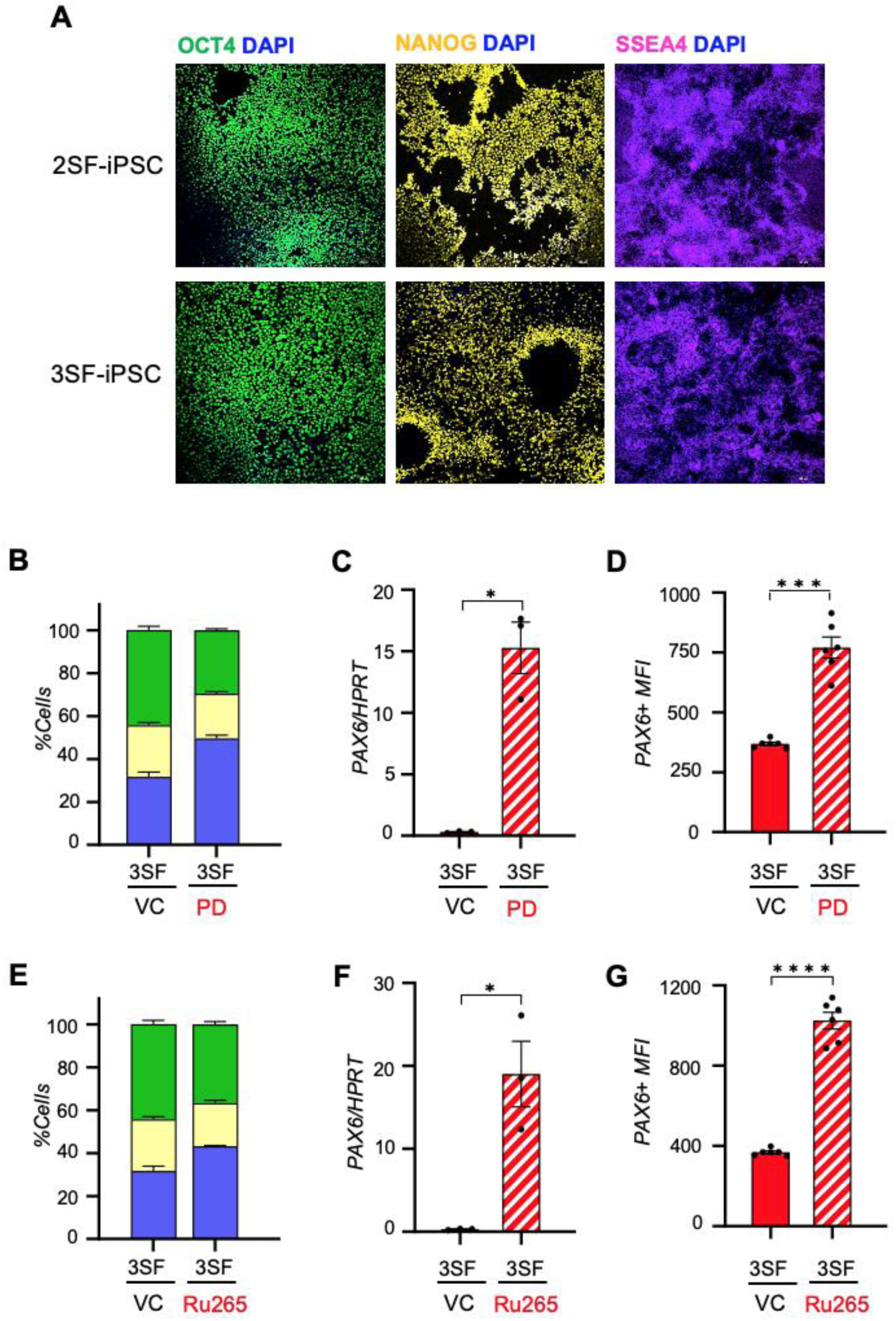
**(A)** Immunohistochemistry for the pluripotency markers OCT4, NANOG and SSEA4 showed uniform staining in the female euploid 2SF and trisomic 3SF iPSC lines. (Scale bar = 50 μm). **(B)** Pretreating the female 3SF-iPSCs for 16 hours with 5µM PD to inhibit cell cycle prior to removal of growth factors to initiate spontaneous differentiation increased the fraction of cells in G_1_/G_0_. **(C)** PD pretreatment of female 3SF-iPSCs increased PAX6 transcript levels and **(D)** PAX6+ median fluorescence intensity on day 5 of spontaneous differentiation. **(E)** Pretreating the female 3SF-iPSCs for 16 hours with 10nM Ru265 to inhibit MCU prior to removal of growth factors to initiate spontaneous differentiation increased the fraction of cells in G_1_/G_0_. **(F)** Ru265 pretreatment of female 3SF-iPSCs increased PAX6 transcript levels and **(G)** PAX6+ median fluorescence intensity on day 5 of spontaneous differentiation. *Transcript levels are normalized to HPRT. Comparisons by Two-tailed T-test. Error bars represent SEM*. *\* P≤0.05, ** P ≤0.01, *** P ≤0.001, **** P ≤0.0001*

**Figure EV6:**
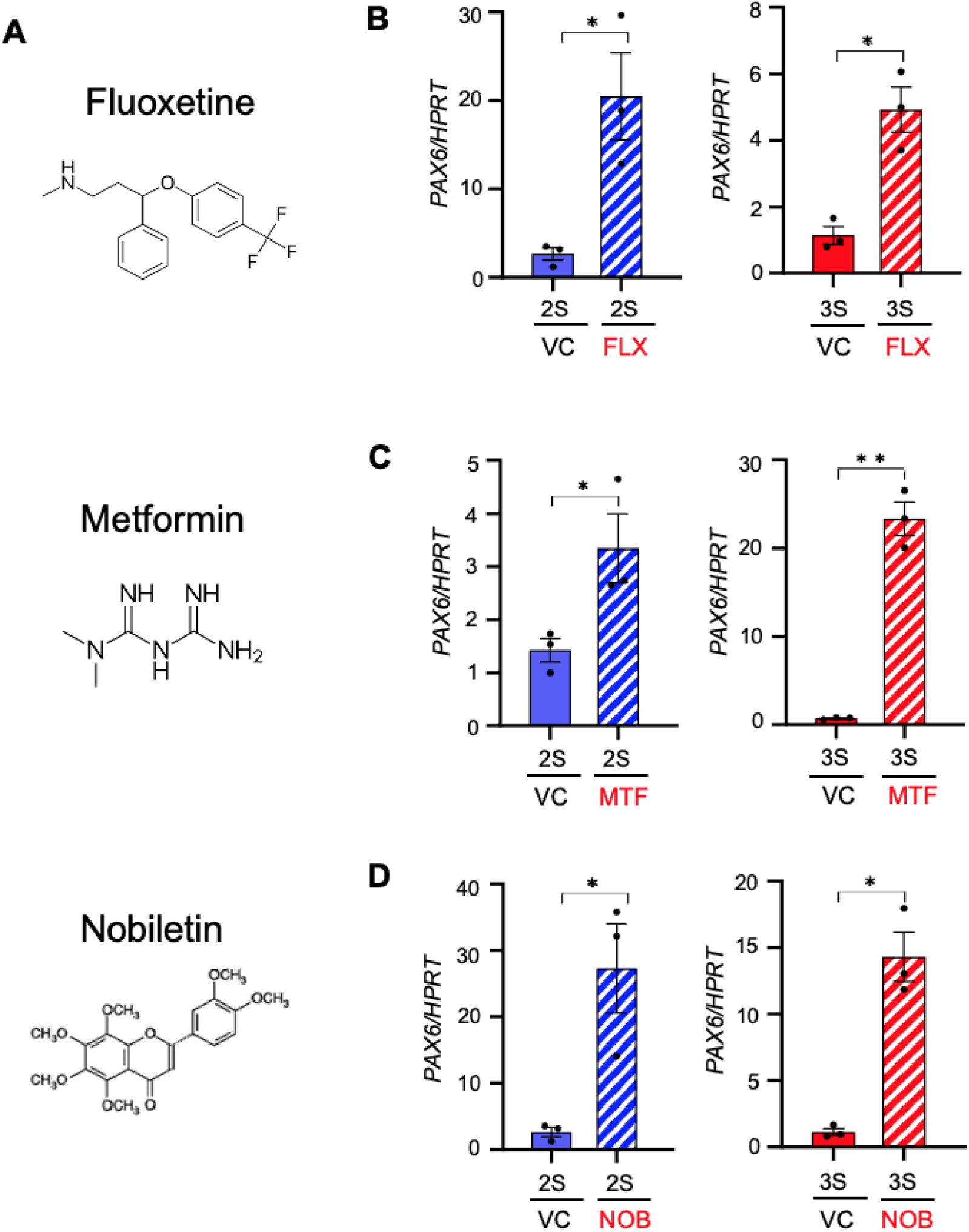
**(A)** Chemical structures for Fluoxetine, Metformin, and Nobiletin. *PAX6* transcript levels on day 5 of spontaneous differentiation increased in both 2S-iPSC and 3S-iPSCs pretreated for 16 hours with (B) 1μM fluoxetine (FLX), (C) 10 mM metformin (MET), or **(D)** 25μM nobiletin (N). *Transcript levels are normalized to HPRT. Comparisons by ANOVA. Error bars represent SEM*. *\* P≤0.05, ** P ≤0.01, *** P ≤0.001, **** P ≤0.0001*

**Figure EV7:**
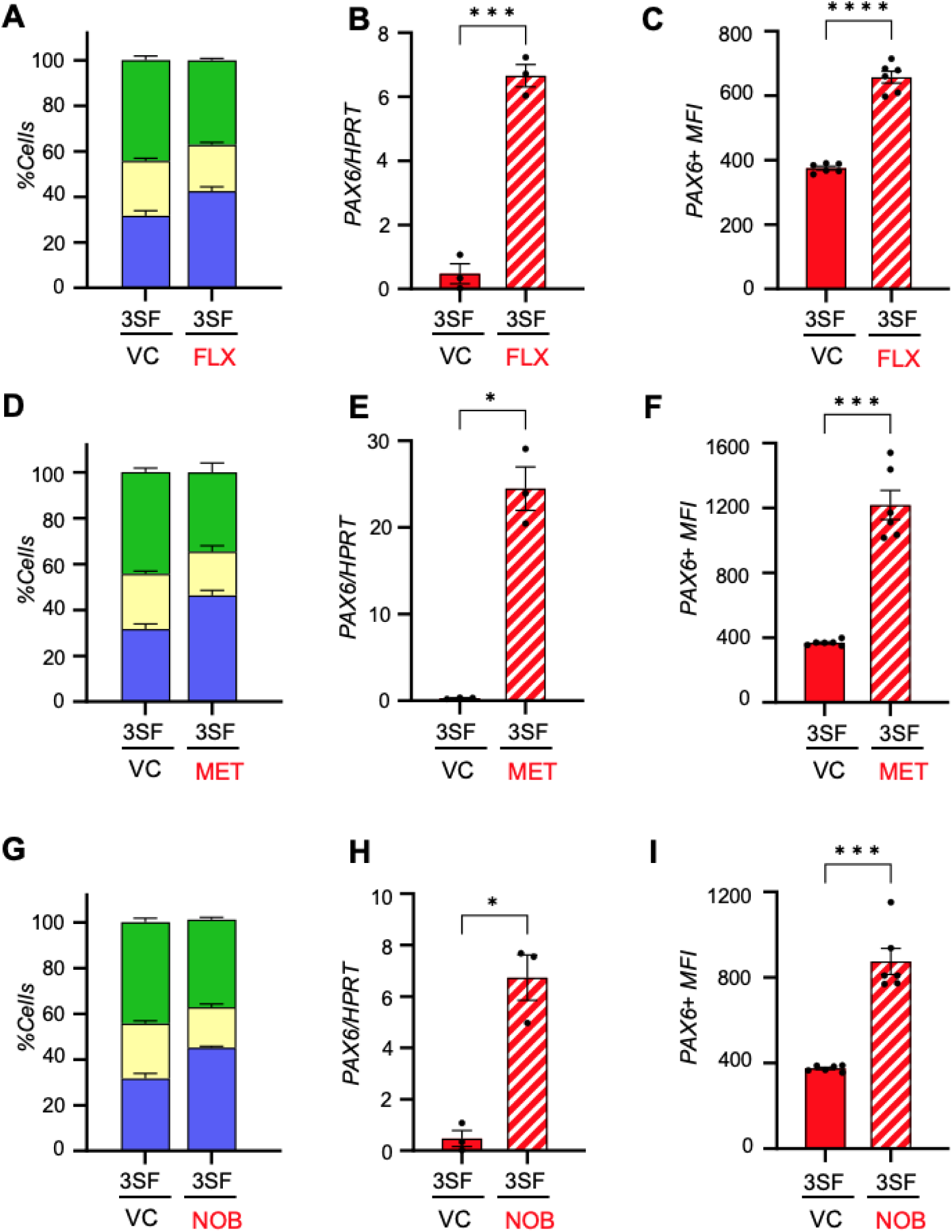
Pretreating the female 3S-iPSCs for 16 hours with 1μM fluoxetine (FLX), 10mM metformin (MET), or 25μM nobiletin (NOB), increased the fraction of cells in G_1_/G_0_ (**A, D, and G**), as well as *PAX6* transcript levels (**B, E and H**) and PAX6+ median fluorescence intensity (**C, F, and I**) on day 5 of spontaneous differentiation. *Transcript levels are normalized to HPRT. Comparisons by ANOVA or Two-tailed T-test. Error bars represent SEM*. *\* P≤0.05, ** P ≤0.01, *** P ≤0.001, **** P ≤0.0001*

**Figure S8:**
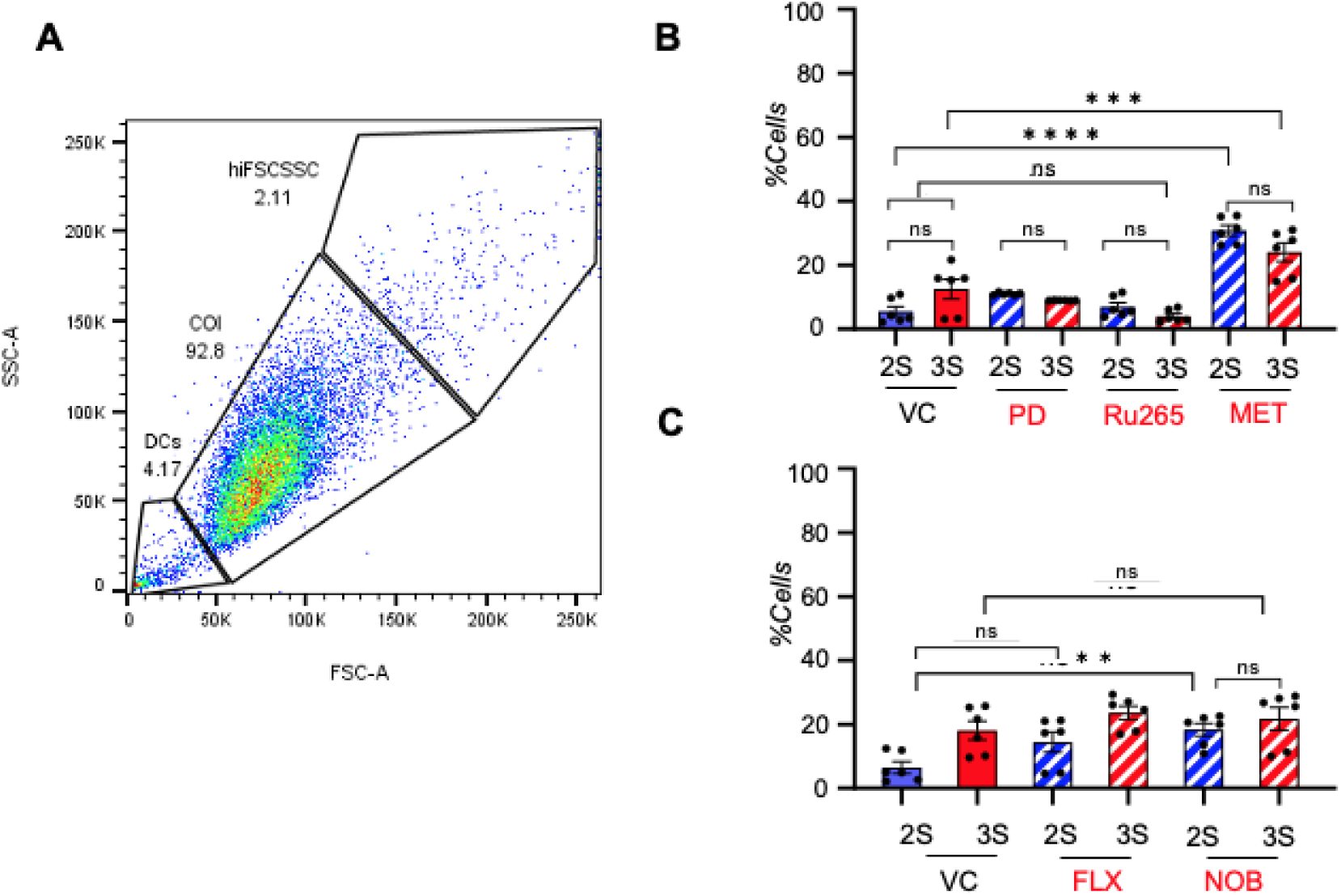
**(A)** Representative gating strategy for selection of cells of interest (COI). Dead cells and debris were eliminated based on a lowFSCSSC signal. Aggregates were eliminated based on a highFSCSSC signal. **(B)** Comparison of the fraction of dead cells after 5 days of differentiation where the vehicle used for drug treatment was water. **(C)** Comparison of the fraction of dead cells after 5 days of differentiation where the vehicle used for drug treatment was DMSO.

**Figure EV9:**
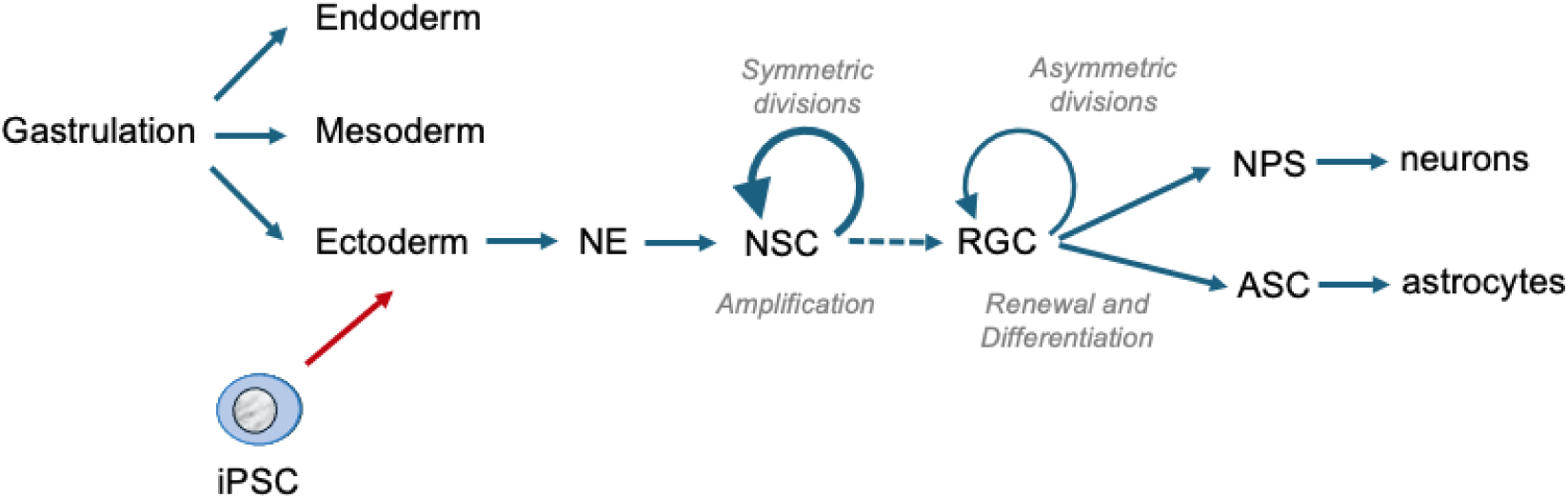
Simplified schematic of cortical neurogenesis. Gastrulation of the embryo establishes the three primary germ cell layers: endoderm, mesoderm, and ectoderm. Ectoderm differentiates to neuroectoderm (NE) which gives rise to neuroepithelial stem cells (NSCs), which ultimately give rise to all components of the CNS. NSCs undergo rounds of symmetrical divisions to expand their own population then gradually transition to radial glial progenitor cells (RGCs), which initiate neurogenesis. RGCs undergo asymmetric divisions which both renew the pool of RGCs and generate either a neural-restricted progenitor cell (NPC) or a glial-restricted progenitor cell, primarily astrocyte progenitor cells (APCs). Spontaneous commitment of iPSCs to NE models one of the earlies steps in this pathway.

**Figure EV10:**
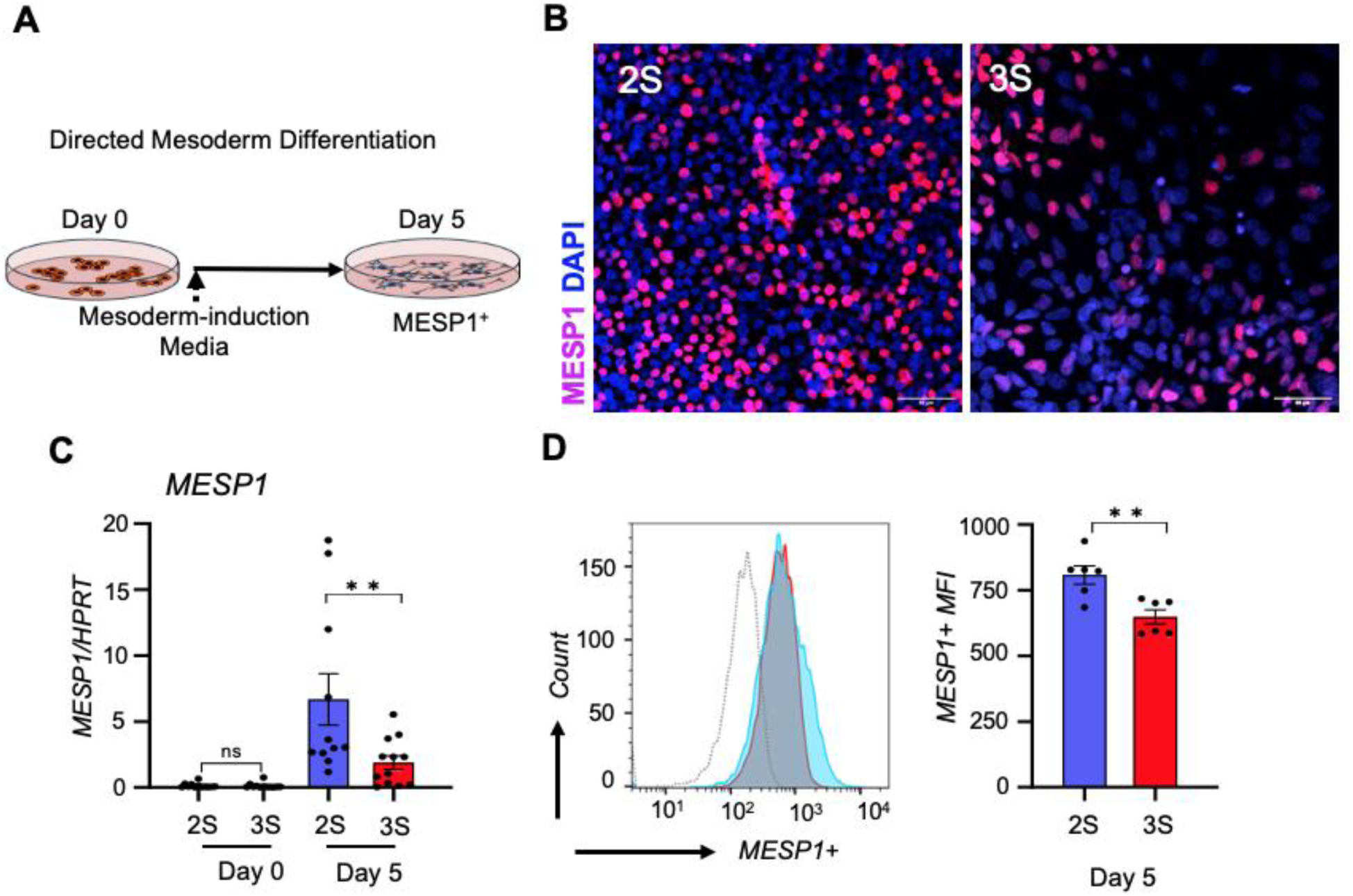
**(A)** Schematic for directed differentiation of iPSCs to Mesoderm **(B)** Fewer cells were MESP1-positive (red) in 3S-derived cultures than in 2S-derived cultures after 5 days of mesoderm-directed differentiation. DAPI-stained nuclei are in blue (Scale bar = 50μm). Both *MESP1* transcript levels **(C)** (n=12) and median MESP1+ intensity **(D)** (n=6) were lower 3S-derived cultures than in 2S-derived cultures on day 5 of directed differentiation. Transcript levels were normalized to *HPRT. Error bars represent SEM.* Significance was determined by ANOVA or a Two-Tailed T-test. *\* P≤0.05, ** P ≤0.01*.

**Figure EV11:**
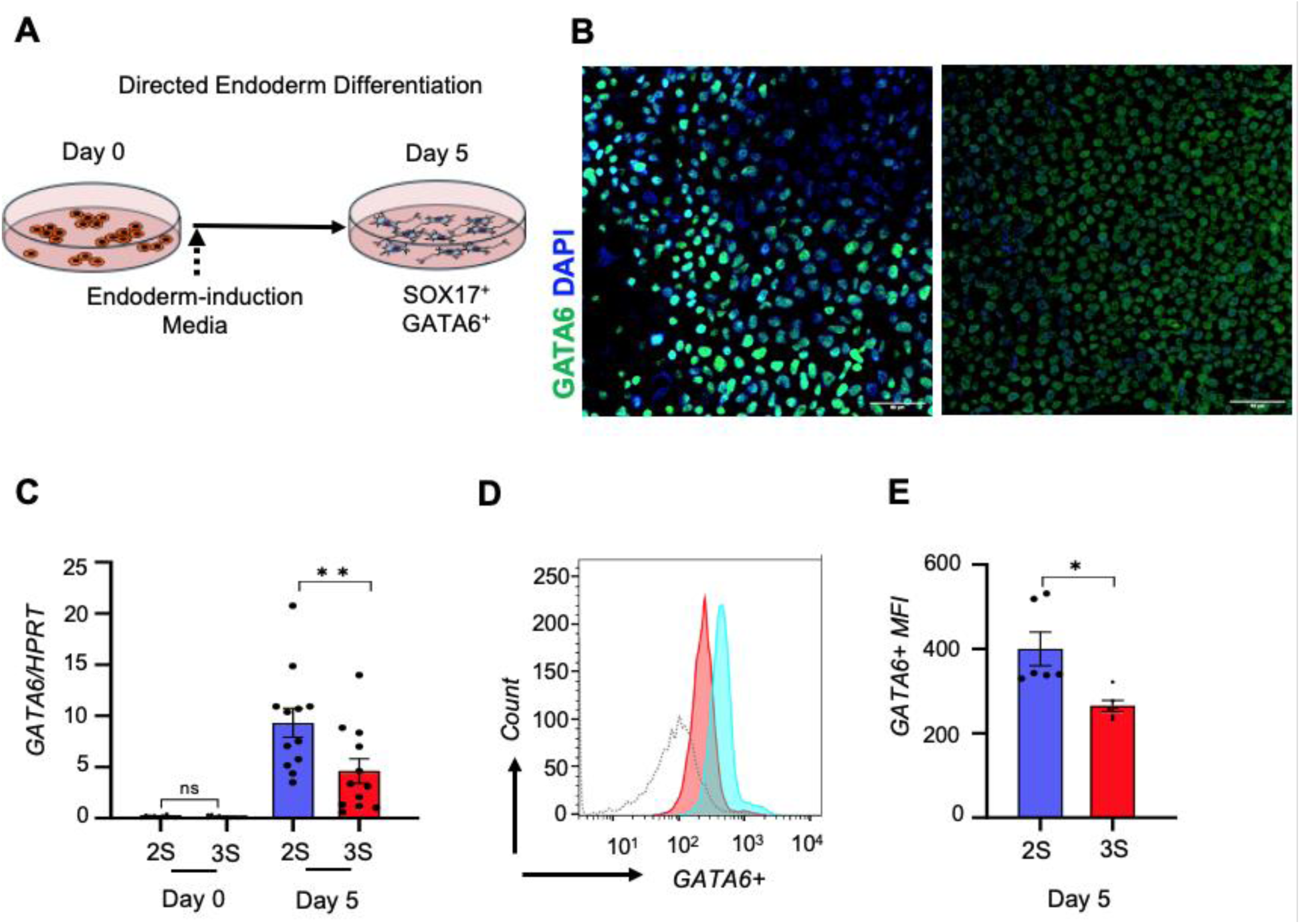
**(A)** Schematic for directed differentiation of iPSCs to endoderm. **(B)** Fewer cells were GATA6-positive (green) in 3S-derived cultures than in 2S-derived cultures after 5 days of endoderm-directed differentiation. DAPI-stained nuclei are in blue (Scale bar = 50μm). Both *GATA6* transcript levels **(C)** (n=12) and median GATA6+ intensity **(D and E)** (n=6) were lower 3S-derived cultures than in 2S-derived cultures on day 5 of directed differentiation. Transcript levels were normalized to *HPRT. Error bars represent SEM.* Significance was determined by ANOVA or Two-Tailed T-test. *\* P≤0.05, ** P ≤0.01*.

